# OxBreaker: species-agnostic pipeline for the analysis of outbreaks using nanopore sequencing

**DOI:** 10.64898/2026.03.18.709804

**Authors:** Carlos Reding, Katie M. V. Hopkins, Matthew Colpus, Nicholas D. Sanderson, Jessica Gentry, Sarah Oakley, Mark Campbell, Drosos Karageorgopoulos, Katie Jeffery, David Eyre, Philip Bejon, Nicole Stoesser, A. Sarah Walker, Bernadette C. Young

## Abstract

Real-time genomic surveillance may mitigate the spread of health-care-associated infections, but whole-genome sequencing costs and the need for specialised expertise constrain its wide implementation in public health. Here we present ‘*OxBreaker*’, an automated and species-agnostic pipeline optimised for the high-resolution analysis of bacterial and plasmid genomes sequenced via Oxford Nanopore Technologies (ONT). ‘*OxBreaker*’ streamlines the transition from raw reads to phylogenetic inference through automated reference selection and high-accuracy variant calling. It is accessible via a graphical user interface (GUI) that can be easily installed locally and operated by non-specialists. Benchmarking against technical and biological replicates of high-priority pathogens demonstrates high accuracy, with false positive variant rates reduced to 0–4 single-nucleotide polymorphisms (SNPs) for common species. We further validated the pipeline by accurately characterising previously published clonal and plasmid-mediated outbreaks, reproducing established phylogenies with improved accessibility. By providing a stable, scalable, open-source offline-compatible solution that matches the resolution of short-read platforms while maintaining the speed of long-read technology, ‘*OxBreaker*’ is designed to facilitate the adoption of local, real-time genomic surveillance for frontline infection prevention and control.

## I. INTRODUCTION

Healthcare-associated infections (HCAIs) represent an important risk to patients and lead to substantial healthcare costs. In England alone, HCAIs affect approximately 8.0%^1^ of acutely hospitalised patients, costing the National Health Service an estimated £2.7 billion annually ^2^. Effective mitigation requires rapid and precise surveillance to differentiate between sporadic cases and active transmission^3^. While genomic surveillance has emerged as a transformative tool, its implementation is frequently hampered by the associated sequencing costs and the specialised bioinformatics expertise required for data interpretation^4,5^.

Short-read sequencing (e.g. Illumina) is widely used but is often centralised due to high capital expenditure and infrastructure requirements^4,6^, leading to turnaround times of days to weeks^7^. Nevertheless, short-read sequencing has led to substantial clinical impact^8^. Furthermore, the inherent limitations of short-read sequencing, with reads 75–300 base pairs (bp) reads in length, impede the resolution of repetitive genomic regions and the tracking of mobile genetic elements (MGEs)^9^, which are major vectors for antimicrobial resistance dissemination in Gram-negative species^10^. The diversity of bacterial genomes^11^ has led to assembly errors with this technology^12^, prompting many clinical studies to default to polymerase chain reaction (PCR)- based multi-locus sequence typing (MLST) and core-genome MLST (cgMLST)^13^. While this approach is robust to assembly errors^13,14^, it lacks the discriminatory power of single-nucleotide polymorphism (SNP)-based analysis required to confirm transmission of highly clonal hospital outbreaks^15,16^.

The alternative approach is long-read sequencing, chiefly through Oxford Nanopore Technologies (ONT). ONT sequencing reads are typically thousands of base pairs long^17^, circumventing some of the issues derived from short-read sequencing. The low entry cost and portability of the MinION platform have the potential to move sequencing from centralised hubs to the hospital bedside. However, the adoption of ONT sequencing for high-resolution outbreak detection has been hindered by higher per-base error rates than short-read sequencing^17^, justifying the use of MLST and cgMLST with ONT platforms^18,19^. Further, a *bioinformatics gap* means clinical laboratories lack the bespoke computational frameworks needed to transform sequence data into a result from which the presence of absence of genomically-linked clusters can be inferred to guide infection prevention and control (IPC).

To bridge this gap, we present ‘*OxBreaker*’, an end-to-end, automated pipeline optimised for the analysis of bacterial genomes sequenced via ONT. ‘*OxBreaker*’, combined with the latest and more accurate ONT chemistry, streamlines the transition from raw reads to phylogenetic inference through an automated routine that encompasses species identification, automated reference selection, and high-accuracy variant calling. Unlike existing tools that rely on continuous internet connectivity, data sharing, and additional costs per sample^20^, ‘*OxBreaker*’ supports an entirely local and offline workflow, ensuring operational resilience against hospital internet outages that could impact diagnostic capabilities in some settings^21^ and data privacy constraints. Importantly, despite being optimised for bacterial genomes, it can also be applied to investigate plasmid transmission.

By distributing ‘*OxBreaker*’ as a conda-compatible package with an integrated Graphical User Interface (GUI), we reduce the need for administrative-level system privileges and command-line proficiency. In other words, our pipeline can be installed and run locally by non-experts, and is robust to software updates of its dependencies thanks to the conda dependency management. We validated the pipeline using two steps: First, through technical and biological replicates of reference strains from the National Collection of Type Cultures (NCTC) representing high-priority HCAIs, and second, via the retrospective analysis of two characterised hospital outbreaks involving both chromosomal and plasmid-mediated transmission. Once validated, we tested ‘*OxBreaker*’ on three new datasets to investigate whether the probability of a common source of transmission between cases in putative outbreaks. Our results demonstrate that ‘*OxBreaker*’ achieves the resolution of short-read platforms while maintaining the speed and accessibility of long-read technology, providing a scalable, local solution for real-time infection prevention and control.

## II. RESULTS

### Pipeline overview

‘*OxBreaker*’ is controlled through a graphical user interface or the command line, depending on whether it is being executed on a local computer or remote cluster. The pipeline requires a path containing FASTQ files with sequencing reads for each isolate. These FASTQ files are generated by `Dorado` during the base-calling process, in our case using the super-accuracy (SUP) algorithm. ONT sequencing generates multiple files per barcode, which means the sequencing reads for one isolate is spread across multiple files and need to be merged into one. ‘*OxBreaker*’ will merge these to provide one FASTQ per barcode—an option that can be disabled to facilitate re-analysis.

Additionally, *OxBreaker*’ requires a reference genome in GenBank format to aid the mapping of the reads to a reference genome. This format includes the genome sequence and the location of coding sequences, including rRNAs and tRNAs used to determine whether a mutation falls into a coding or non-coding regions. Here the user can provide the reference genome, but if none is provided—for example because the species investigated is unknown—‘*OxBreaker*’ can find the closest reference genome available in the Reference Sequence (RefSeq) database. Finally, there are options for the user to provide a random seed for reproducibility, and run the pipeline. Once the user clicks on ‘Run Pipeline’, ‘*OxBreaker*’ will run through a set of stages illustrated in Figure 1.

**Figure 1.**
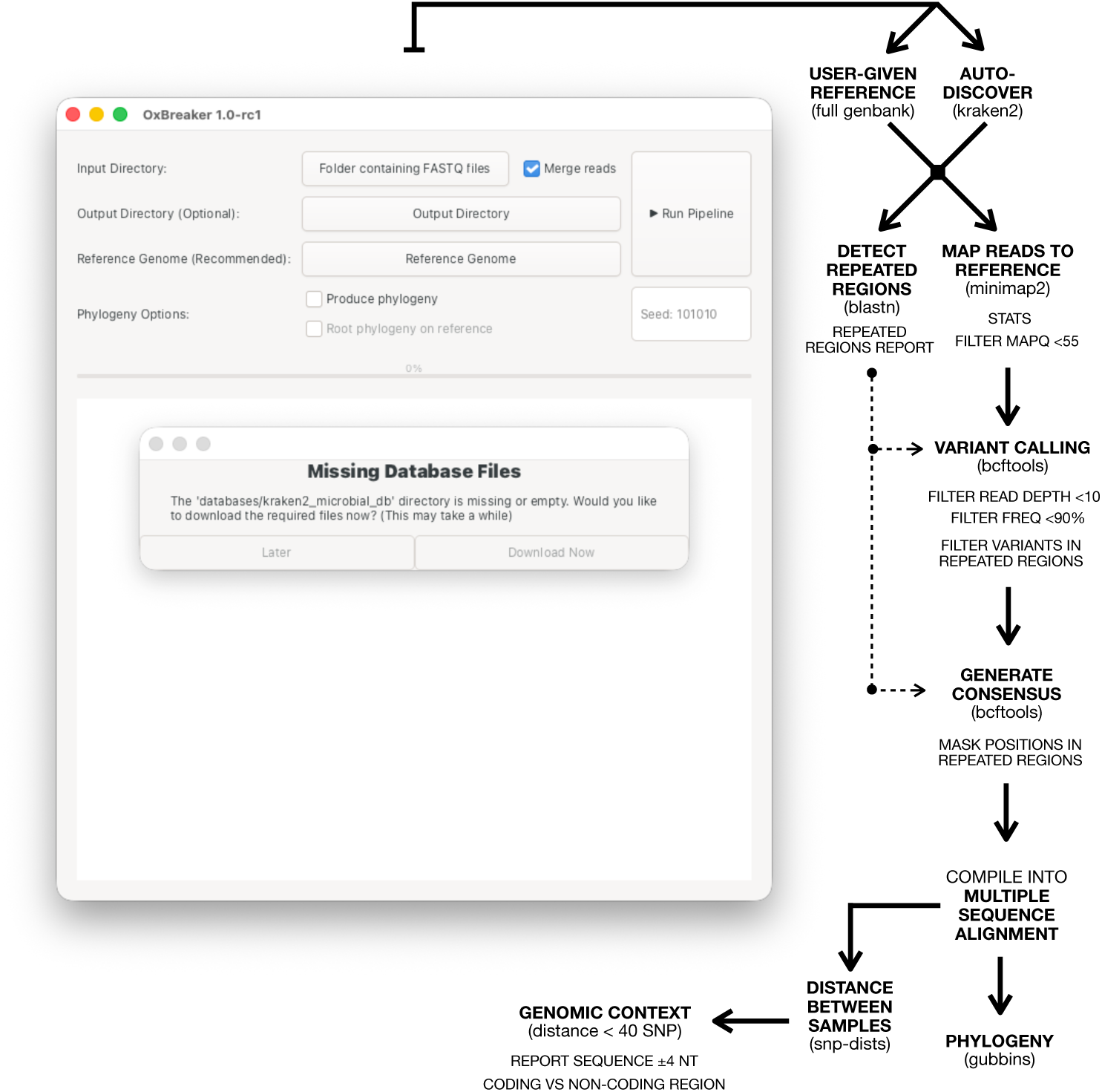
User interface and pipeline overview. The graphical user interface (left) allows the user to specify the location of the reads as compressed FASTQ files (required), whether read files need merging to produce one file per barcode (default: no barcode merging), where to store the results (default: reads’ parent directory), and the GenBank file containing sequence and gene location metadata used as reference (default: auto-discover). It also contains a button to specify whether to use the reference genome to root the phylogeny, and another run the pipeline alongside a small terminal showing the progress. Once the *run* button is pressed, the interface spawns nextflow to run the pipeline (right). In the simplest scenario, a user would have to specify where the FASTQ files are located, and press *run*. ‘*OxBreaker*’ will prompt the user to download the Kraken2 Microbial Database (30GB) when run for the first time, if it is not detected in the system.

‘*OxBreaker*’ pools all samples into a temporary file that is run through `Kraken2`^22^ using the 30GB microbial database^23^ when a reference genome is not given. It then sorts the species and specific strains reported by number of reads assigned to them and programmatically extracts information about references from the NCBI’s RefSeq database^24^ using `biopython`^25^. The workflow prioritises complete and resolved reference genomes by checking the candidate reference’s metadata to verify it only contains one contig. If the candidate contains multiple contigs, ‘*OxBreaker*’ will check whether the assembly files contain plasmids and shortlist the candidate only if the chromosome is resolved into one contig. Otherwise, the next most frequently reported database match is checked. If none of the candidate strains meet these requirements, the run is halted with a warning. In this case, the pipeline will stop and ask the user for a reference genome. Once the best match is found, ‘*OxBreaker*’ downloads the reference sequence. Since GenBank files are not part of the NCBI’s assembly database, ‘*OxBreaker*’ retrieves the sequence in FASTA format and populates the accession code in the header to retrieve the full GenBank report from the NCBI’s nucleotide database, containing sequence and gene location metadata.

‘*OxBreaker*’ then analyses the reference genome to detect and mask repeated regions and avoid false positives introduced by `Dorado` during base-calling of repeated sequences^26–28^. This uses blastn from the BLAST+ suite^29^ with default settings, meaning these regions must be at least 11nt long, with the reference acting as both reference and query sequences^30,31^. ‘*OxBreaker*’ stores the location of sequences that are found more than once and produces a list of coordinates containing non-repeated sequences in BED format.

The sequence reads, pooled into one file per barcode, are then mapped to the reference genome using minimap2^32^ version 2.29-r1283, with settings for high-quality long-read data (`-x lr:hq`), outputting a file in SAM format. Additional options `-a --secondary=no --sam-hit--only` are specified to ignore secondary alignments and reads that were not mapped to the reference. Importantly, ‘*OxBreaker*’ uses a seed in `minimap2` to ensure reproducibility of this step which can be set by the user with the option `--seed 12345` or through the user interface.

The resulting filtered alignment is then fed to `bcftools`^33^ version 1.23 for variant calling. ‘*OxBreaker*’ uses the new multiallelic caller (`--multiallelic-caller`), and passes through the previously provided BED file so `bcftools` ignores coordinates that correspond to repeated sequences in the reference genome. Quality checks, including minimum supporting read depth, minimum allele frequency, masking variant calls in repeat regions, and exclusion of small insertion-deletions are applied when ‘*OxBreaker*’ post-processes the variants.

To calculate the distance between potentially related isolates, ‘*OxBreaker*’ generates a consensus for each isolate by mapping the variants found in that isolate onto the reference genome used with `bcftools`. Then, the consensus sequences are compiled into a multiple sequence alignment file which ‘*OxBreaker*’ feeds to `snp`-`dists`^34^ to compute the distance between the samples and produce a phylogeny as a final step with `gubbins`^35^. To root the phylogeny to the reference, the user can pass `--debug` to the command line. SNP thresholds for outbreak detection vary across different species and technologies, from 0–2 SNPs^36^ to 40 SNPs^37,38^. In this species agnostic pipeline, and particularly given the potential for incomplete sampling of all cases within a transmission chain, possible genomic links are reported if up to 40 variants are identified between two sequences. In this case ‘*OxBreaker*’, reports the genomic context for each variant, 4nt up- and downstream of the variant to retrieve its genomic context, as well as whether variants are located in non-coding or coding regions—the latter suggesting a mutation might be associated with a phenotype change.

### First calibration step: Filtering by mapping quality minimises data loss

To calibrate quality control and other pipeline parameters, we repeatedly sequenced NCTC isolates from a range of Gram-negative and Gram-positive pathogens implicated in HCAIs for which reference sequence data were also available (denoted the ‘replicate reference set’): *Clostridioides difficile* 630, *C. difficile* R20291, *Escherichia coli* CFT073, *Klebsiella pneumoniae* MGH 78578, *Pseudomonas aeruginosa* PA01, and *Staphylococcus aureus* MRSA252 (Table 1).

**Table 1.** Summary of the replicate reference set. Detail of all species and replicates analysed, with depths calculated from mapping without read quality filtering in Figure S1.

| Species | Biological*<br>Replicates | Technical*<br>Replicates | Biological<br>Replicate Depth | Technical<br>Replicate Depth | Source |
| --- | --- | --- | --- | --- | --- |
| *C. difficile* 630 | 2 | 1 | 12<sup>†</sup>/46 | 28 | This study |
| *C. difficile* R20291 | 2 | 1 | 22/18 | 7<sup>†</sup> | This study |
| *E. coli* CFT073 | 2 | 1 | 81/108 | 99 | This study and [43] |
| *K. pneumoniae* MGH 78578 | 2 | 1 | 91/270 | 244 | This study and [43] |
| *P. aeruginosa* PA01 | 2 | 1 | 60/95 | 79 | This study and [43] |
| *S. aureus* MRSA252 | 2 | 1 | 29/867 | 627 | This study and [43] |

\* ‘Biological replicate’ means the isolate was grown separately before sequencing. ‘Technical replicate’ means that DNA was extracted from one biological replicate and divided into two before sequencing. Technical replicates are always paired: the sequence with the greatest depth was classified as the biological replicate, and the lower depth as the technical replicate.
<sup>†</sup> Less than 75% of the reference genome available for analysis and thus excluded by ‘OxBreaker’.

As expected, given the strain-specific inconsistencies in ONT reads sequencing^39^, yields varied within, as well as across, flow cells. However, sequencing yields of individual flow cells varied substantially, leading to coverage ranging from 7 to 867-fold before read quality filtering despite loading the same amount of DNA (Table 1). Considering read quality-filtering as the first possible quality-control step, filtering reads with less than 99.9% accuracy (<Q30) resulted in a loss of 95.1%±3.9% (mean ± standard deviation) of depth across all replicates (Figures 3 and S1). Relaxing the quality threshold to remove reads with less than 99% accuracy (<Q20) mitigated the loss of data, but still 30.5%±14.3% of depth was lost across the replicate reference set. Loss of data depth was significant even when filtering reads with an accuracy of ∼60% (<Q4, 2-sided Wilcoxon signed-rank p=0.0015). However, this loss in depth did not necessarily translate into a proportional loss in breadth of coverage of the reference genome, at least at depth 10 (Figure S2).

We investigated false positive variant detection and genome coverage when reads were filtered across a range of quality thresholds, and found no evidence that, given the mapping quality filtering described above, additional filtering based on read quality affected variants called. However, it did substantially reduce the percentage of the genome called after mapping (Figure S3). Thus, we did not include additional read filtering based on quality.

Next, we explored filtering the data based the mapping of the reads. As Figure 2A illustrates, 96.9% of all reads from the pooled replicate reference set sequence reads mapped almost perfectly to the reference sequence (mapping quality score = 60, only 0.000001% probability that the sequence did not originate from that region in the reference). Sequences from real, potential outbreak isolates can be more diverse, with samples being more distantly related to any single reference. For example, mapping a second dataset containing a potential outbreak of *C. difficile* isolates to the genome of *C. difficile* 630, and the mean number of reads mapped almost perfectly to the reference was lower (88.9%±16.3%, Figure 2B). Thus, to account for such diversity, ‘*OxBreaker*’ filters the resulting alignment to remove reads with a mapping quality <55— approximately three times the error of a mapping quality score of 60—and sorts the result by coordinates using `samtools`^33^.

**Figure 2.**
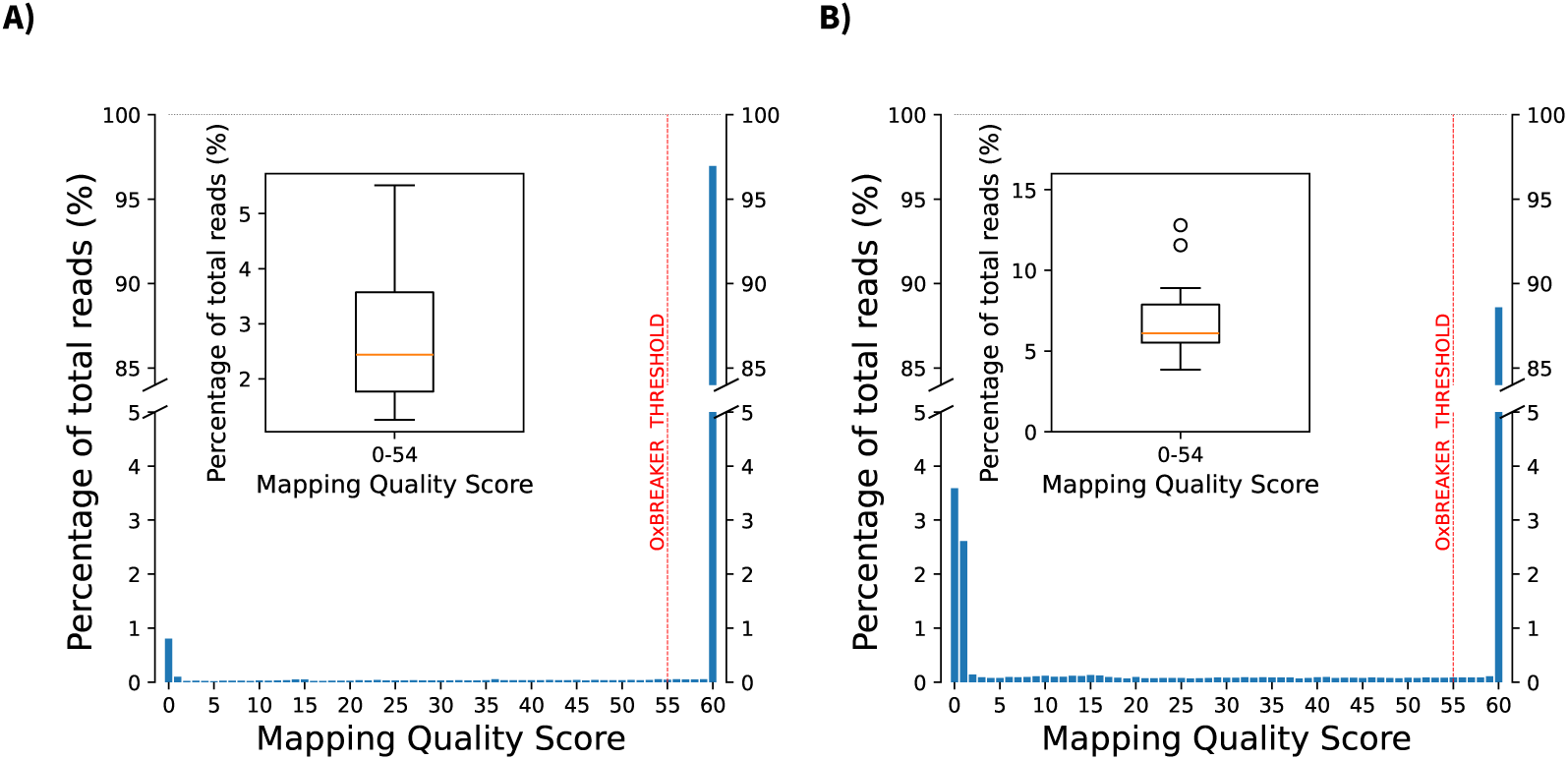
Mapping quality scores in the replicate reference set (A) and a diverse potential *C. difficile* outbreak (B). Distribution of quality scores pooled across all strains analysed. The distribution is bi-modal with a mode at score 60 meaning perfect match to its corresponding reference sequence, comprising 96.9% of all reads for the replicate reference set (A) and 88.9% for the diversified *C. difficile* set (B), and another at score 0, meaning reads mapping to multiple locations, comprising 0.9% and 11.1% of all reads, respectively. The red, vertical line corresponds to the minimum quality score imposed by ‘*OxBreaker*’ (55) and includes 97.1%±1.3% of all reads in the replicate reference set (A), whereas for the diversified *C. difficile* (B) it includes 88.9±16.3% of all reads. The insets show the percentage of reads with a mapping score below 55. The red line represents the median, the box is the inter-quartile range (IQR, distance between 25th and 75th percentiles), and the whiskers represent data corresponding to 1.5×IQR above and below the 75th and 25th percentiles, respectively.

**Figure 3.**
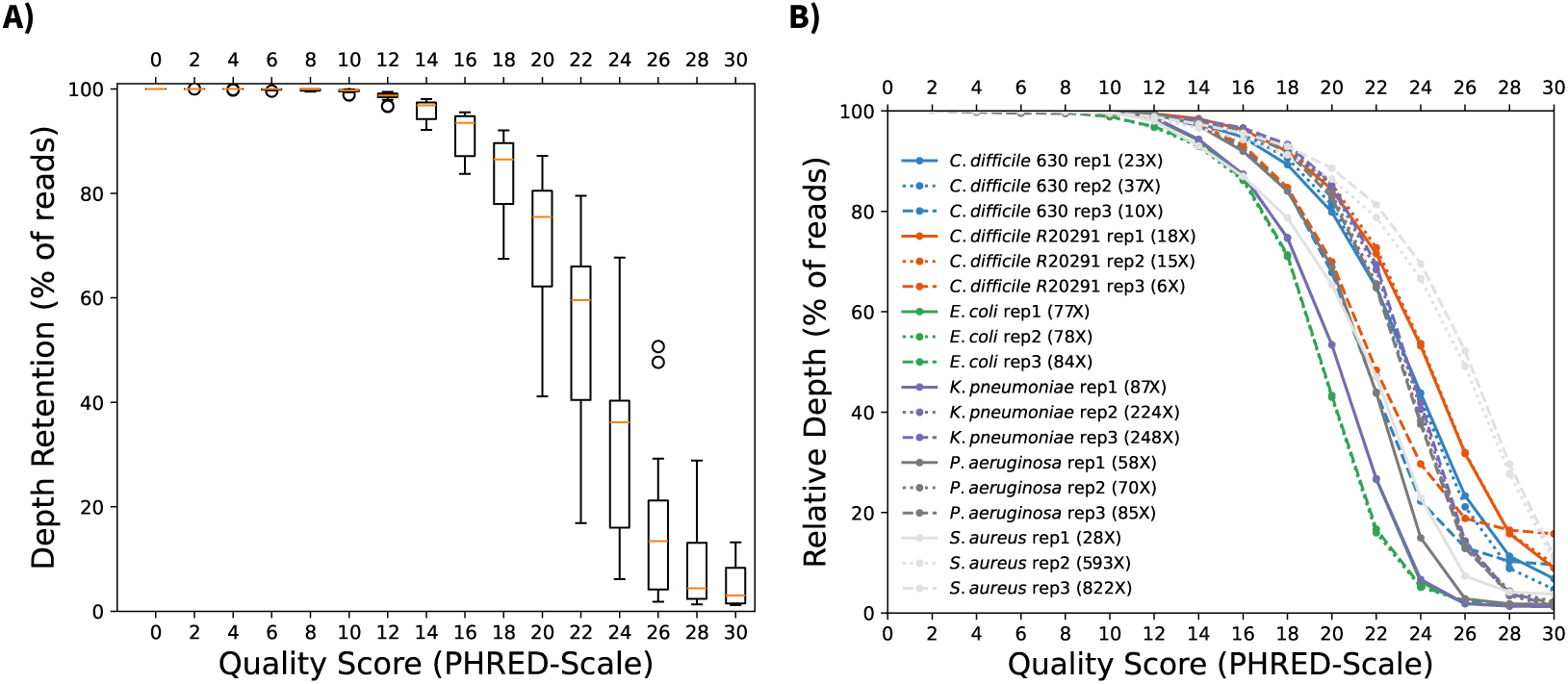
Depth loss resulting from filtering reads by quality in the replicate reference set. **A)** Percentage of the reads retained across the replicate reference set. Depth data was normalised to the mean depth per replicate, without quality filtering (Q0), to calculate the relative depth lost when filtering reads below a given quality score ranging. The red line represents the median, the box is the inter-quartile range (IQR, distance between 25th and 75th percentiles), and the whiskers represent data corresponding to 1.5 × IQR above and below the 75th and 25th percentiles, respectively. Outliers are represented as dots. We ran a 2-sided Wilcoxon signed-rank test to compare the loss of depth without filtering (Q0), and after filtering reads below Q4 (*p* ≈ 0.0015), Q6 (*p <* 10^−4^) and Q8 (*p <* 10^−4^)—no read had a score below 2, resulting in an identical dataset to no filtering. **B)** Normalised depth of each individual sample after filtering reads with quality scores below 0–30. The number shown in the legend is the absolute depth for each sample with no read quality filtering.

### Second calibration step: Filtering variants by supporting depth and frequency minimises false positives

The sequences from the replicate reference set (Table 1) should exactly match their corresponding true reference sequences, and thus should have 0 variants compared with the reference. Any variants detected are therefore by definition false positives. After filtering the reads by mapping quality score, we investigated the impact of filtering positions based on their depth and the frequency (percentage) of reads supporting a variant comprised of at least that read depth, for combinations of read depth from 1 to 30, and percentage frequencies from >0% to 100% (Figure 4, coverage in Figure S4, extended data for isolates with grater coverage in Figure S5). Despite the differences in sequencing depth across the sequenced reference isolates, we found that read depth had a much smaller effect on false positives than expected, restricted mostly to cases with no or very relaxed frequency constraints. In contrast frequency had a greater effect, with 90% frequency reducing false positives compared with 80%, and with relatively little difference versus 100% across species. Overall, a minimum supporting depth of 10, to accommodate variability in sequencing yields, and a minimum variant frequency of 90%, reduced the number of false positives versus the original true reference to 0–4 in both strains of *C. difficile*, *E. coli*, and *K. pneumoniae* (Table 2).

**Figure 4.**
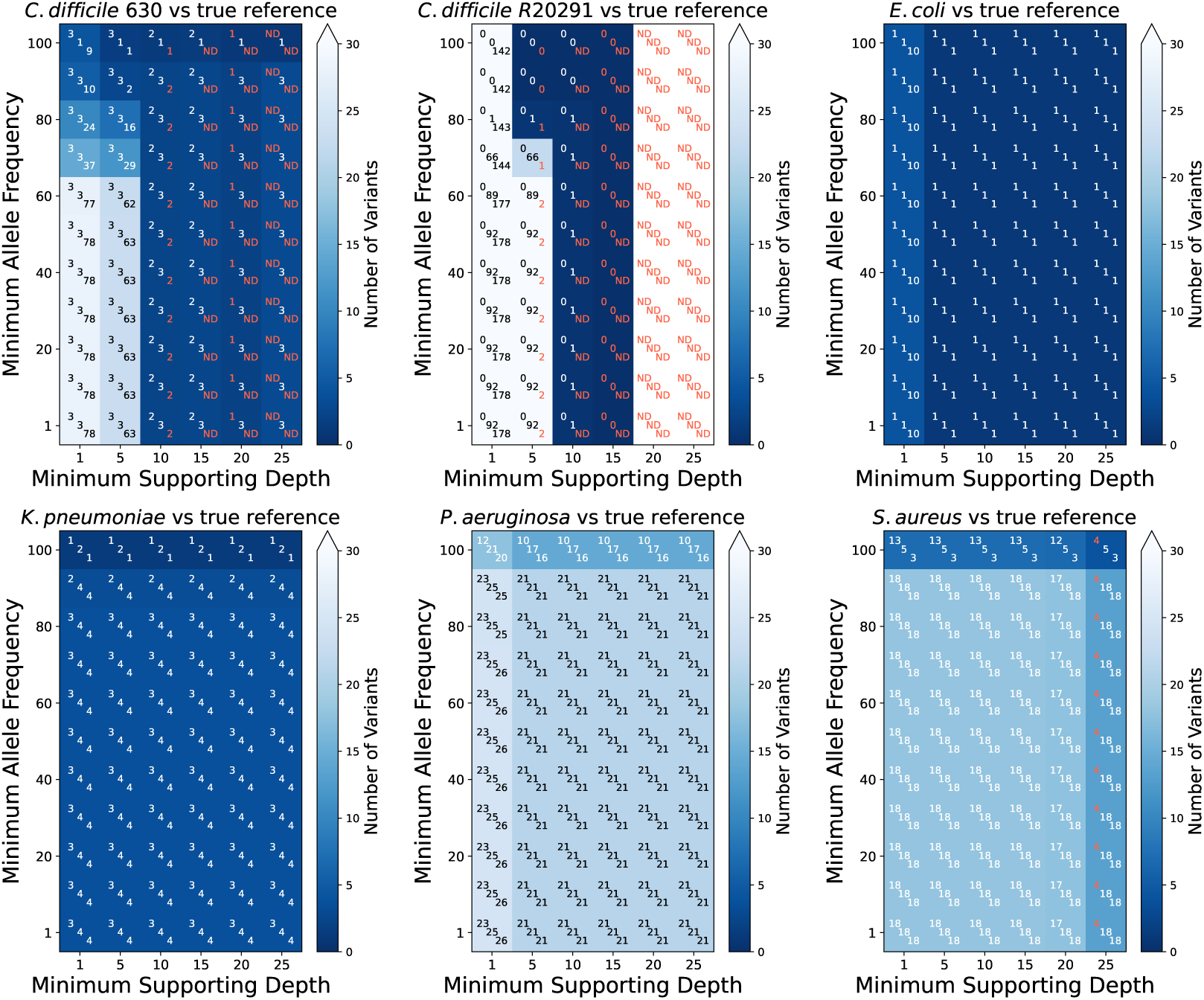
Number of false positives, compared with the true reference, in the replicate reference set by minimum frequency and read depth supporting each call. The number of false positives for each replicate is shown in the centre of each cell, comparing each replicate with the reference genome: Cells coloured based on the number of false positives found with darker tones representing fewer differences (theoretical minimum 0). Red text indicates the number of variants found in combinations where less than 75% of the reference genome was called, due to the loss of breadth of coverage resulting from increasing the supporting depth (Figure S2). This suggests that the fewer variants detected is likely caused by loss of data rather than optimal combination of variant frequency and supporting depths. Samples with less than the required depth are indicated by ND (no data). Figure S4 shows this coverage and Figure S5 shows extended data with higher sequencing depths.

**Table 2.**
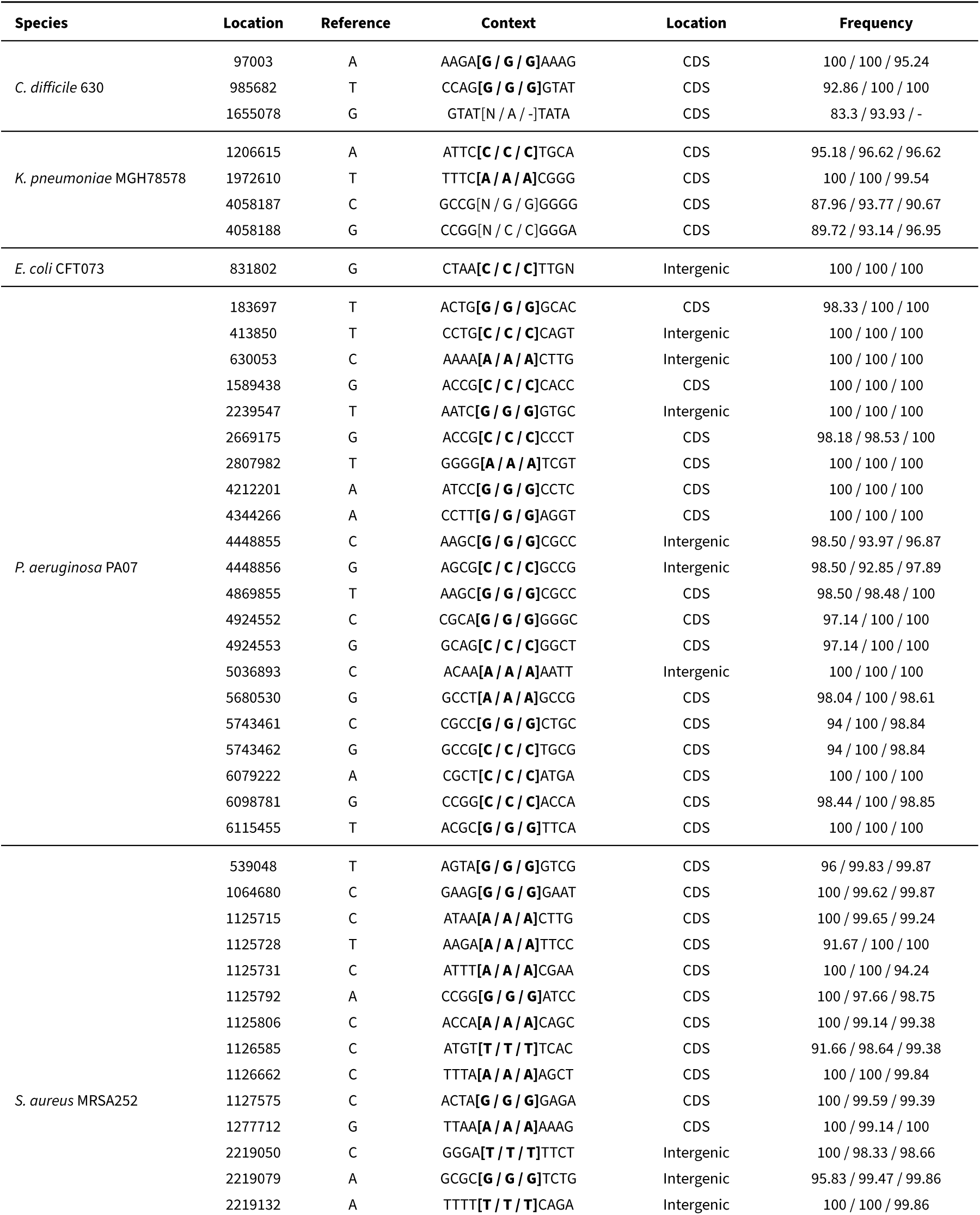
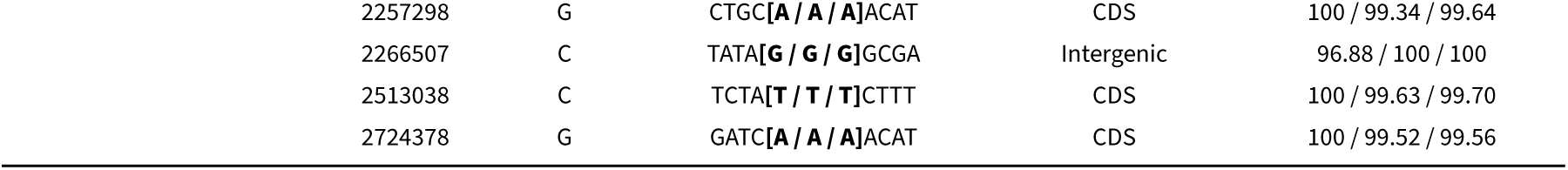
Summary of false positive variants in the replicate reference set. For each location with any variant called in any isolate in the replicate reference set (required to have at least 90% of frequency and a depth of 10), ‘context’ shows the 4 nucleotides before and after the variant position, then the square brackets show the nucleotides called in the replicates, ordered from highest to lowest coverage. Bold underline text indicates the same variant called in all three replicates, potentially indicating a change in the biological isolate compared with the original reference strain. *CDS* = conding sequence, *-* = replicate excluded by ‘*OxBreaker*’ due to sample having less than 75% of the genome mapped to the reference.

| Species | Location | Reference | Context | Location | Frequency |
| --- | --- | --- | --- | --- | --- |
| *C. difficile* 630 | 97003 | A | AAGA[G / G / G]AAAG | CDS | 100 / 100 / 95.24 |
|  | 985682 | T | CCAG[G / G / G]GTAT | CDS | 92.86 / 100 / 100 |
|  | 1655078 | G | GTAT[N / A / -]TATA | CDS | 83.3 / 93.93 / - |
| *K. pneumoniae* MGH78578 | 1206615 | A | ATTC[C / C / C]TGCA | CDS | 95.18 / 96.62 / 96.62 |
|  | 1972610 | T | TTTC[A / A / A]CGGG | CDS | 100 / 100 / 99.54 |
|  | 4058187 | C | GCCG[N / G / G]GGGG | CDS | 87.96 / 93.77 / 90.67 |
|  | 4058188 | G | CCGG[N / C / C]GGGA | CDS | 89.72 / 93.14 / 96.95 |
| *E. coli* CFT073 | 831802 | G | CTAA[C / C / C]TTGN | Intergenic | 100 / 100 / 100 |
| *P. aeruginosa* PA07 | 183697 | T | ACTG[G / G / G]GCAC | CDS | 98.33 / 100 / 100 |
|  | 413850 | T | CCTG[C / C / C]CAGT | Intergenic | 100 / 100 / 100 |
|  | 630053 | C | AAAA[A / A / A]CTTG | Intergenic | 100 / 100 / 100 |
|  | 1589438 | G | ACCG[C / C / C]CACC | CDS | 100 / 100 / 100 |
|  | 2239547 | T | AATC[G / G / G]GTGC | Intergenic | 100 / 100 / 100 |
|  | 2669175 | G | ACCG[C / C / C]CCCT | CDS | 98.18 / 98.53 / 100 |
|  | 2807982 | T | GGGG[A / A / A]TCGT | CDS | 100 / 100 / 100 |
|  | 4212201 | A | ATCC[G / G / G]CCTC | CDS | 100 / 100 / 100 |
|  | 4344266 | A | CCTT[G / G / G]AGGT | CDS | 100 / 100 / 100 |
|  | 4448855 | C | AAGC[G / G / G]CGCC | Intergenic | 98.50 / 93.97 / 96.87 |
|  | 4448856 | G | AGCG[C / C / C]GCCG | Intergenic | 98.50 / 92.85 / 97.89 |
|  | 4869855 | T | AAGC[G / G / G]CGCC | CDS | 98.50 / 98.48 / 100 |
|  | 4924552 | C | CGCA[G / G / G]GGGC | CDS | 97.14 / 100 / 100 |
|  | 4924553 | G | GCAG[C / C / C]GGCT | CDS | 97.14 / 100 / 100 |
|  | 5036893 | C | ACAA[A / A / A]AATT | Intergenic | 100 / 100 / 100 |
|  | 5680530 | G | GCCT[A / A / A]GCCG | CDS | 98.04 / 100 / 98.61 |
|  | 5743461 | C | CGCC[G / G / G]CTGC | CDS | 94 / 100 / 98.84 |
|  | 5743462 | G | GCCG[C / C / C]TGCG | CDS | 94 / 100 / 98.84 |
|  | 6079222 | A | CGCT[C / C / C]ATGA | CDS | 100 / 100 / 100 |
|  | 6098781 | G | CCGG[C / C / C]ACCA | CDS | 98.44 / 100 / 98.85 |
|  | 6115455 | T | ACGC[G / G / G]TTCA | CDS | 100 / 100 / 100 |
| *S. aureus* MRSA252 | 539048 | T | AGTA[G / G / G]GTCG | CDS | 96 / 99.83 / 99.87 |
|  | 1064680 | C | GAAG[G / G / G]GAAT | CDS | 100 / 99.62 / 99.87 |
|  | 1125715 | C | ATAA[A / A / A]CTTG | CDS | 100 / 99.65 / 99.24 |
|  | 1125728 | T | AAGA[A / A / A]TTCC | CDS | 91.67 / 100 / 100 |
|  | 1125731 | C | ATTT[A / A / A]CGAA | CDS | 100 / 100 / 94.24 |
|  | 1125792 | A | CCGG[G / G / G]ATCC | CDS | 100 / 97.66 / 98.75 |
|  | 1125806 | C | ACCA[A / A / A]CAGC | CDS | 100 / 99.14 / 99.38 |
|  | 1126585 | C | ATGT[T / T / T]TCAC | CDS | 91.66 / 98.64 / 99.38 |
|  | 1126662 | C | TTTA[A / A / A]AGCT | CDS | 100 / 100 / 99.84 |
|  | 1127575 | C | ACTA[G / G / G]GAGA | CDS | 100 / 99.59 / 99.39 |
|  | 1277712 | G | TTAA[A / A / A]AAAG | CDS | 100 / 99.14 / 100 |
|  | 2219050 | C | GGGA[T / T / T]TTCT | Intergenic | 100 / 98.33 / 98.66 |
|  | 2219079 | A | GCGC[G / G / G]TCTG | Intergenic | 95.83 / 99.47 / 99.86 |
|  | 2219132 | A | TTTT[T / T / T]CAGA | Intergenic | 100 / 100 / 99.86 |

|  |  |  |  |  |
| --- | --- | --- | --- | --- |
| 2257298 | G | CTGC[**A / A / A**]ACAT | CDS | 100 / 99.34 / 99.64 |
| 2266507 | C | TATA[**G / G / G**]GCGA | Intergenic | 96.88 / 100 / 100 |
| 2513038 | C | TCTA[**T / T / T**]CTTT | CDS | 100 / 99.63 / 99.70 |
| 2724378 | G | GATC[**A / A / A**]ACAT | CDS | 100 / 99.52 / 99.56 |

For *P. aeruginosa* and *S. aureus*, however, ‘*OxBreaker*’ found 21 and 18 false positives respectively at these thresholds. In all cases the false positives were either shared among all replicates of each species or the same variant was called in all replicates but did not pass the minimum depth/frequency requirements imposed by ‘*OxBreaker*’. This was also the case for the smaller number of false positives found in other species. This suggests mutations could have accumulated in the local copies of the reference isolates following multiple laboratory passages over the years. Consequently, comparing replicates with each other, they were all either identical at variant sites or variant positions were masked and not taken into account in the pairwise comparisons from `snp`-`dists` (Table 2, Figure S6). Of note, a previous implementation of ‘*OxBreaker*’ implementing Clair3^40^ for variant-calling failed this control, with different false positives between replicates of the same isolate when mapped to the true reference (Figure S6 and S7) and conflicting with the contents of the read alignment files—contradicting the reported superiority of deep learning variant callers on bacterial ONT sequence data^41^.

To test the robustness of these thresholds, we repeated the analysis above using a reference genome of the same species that was phylogenetically much less related to the isolates analysed. For *C. difficile* 630 we used *C. difficile* R20291, for *E. coli* CFT073 *E. coli* K-12 MG1655 (accession code NC_000913), for *K. pneumoniae* MGH78578 *K. pneumoniae* KP4831 (NZ_CP09445), for *P. aeruginosa* PA01 *P. aeruginosa* B136-33 (NC_020912), and for *S. aureus* MRSA252 *S. aureus* MSSA ST15 (CP012972). While the number of variants found with respect to the reference increased, as expected, we found a range of depth and frequency combinations with 0–1 variants with respect to the replicates of the same isolate that were concordant with our thresholds (Figure 5) in terms of false positive discovery. In other words, introducing a distant reference did not introduce false positives.

**Figure 5.**
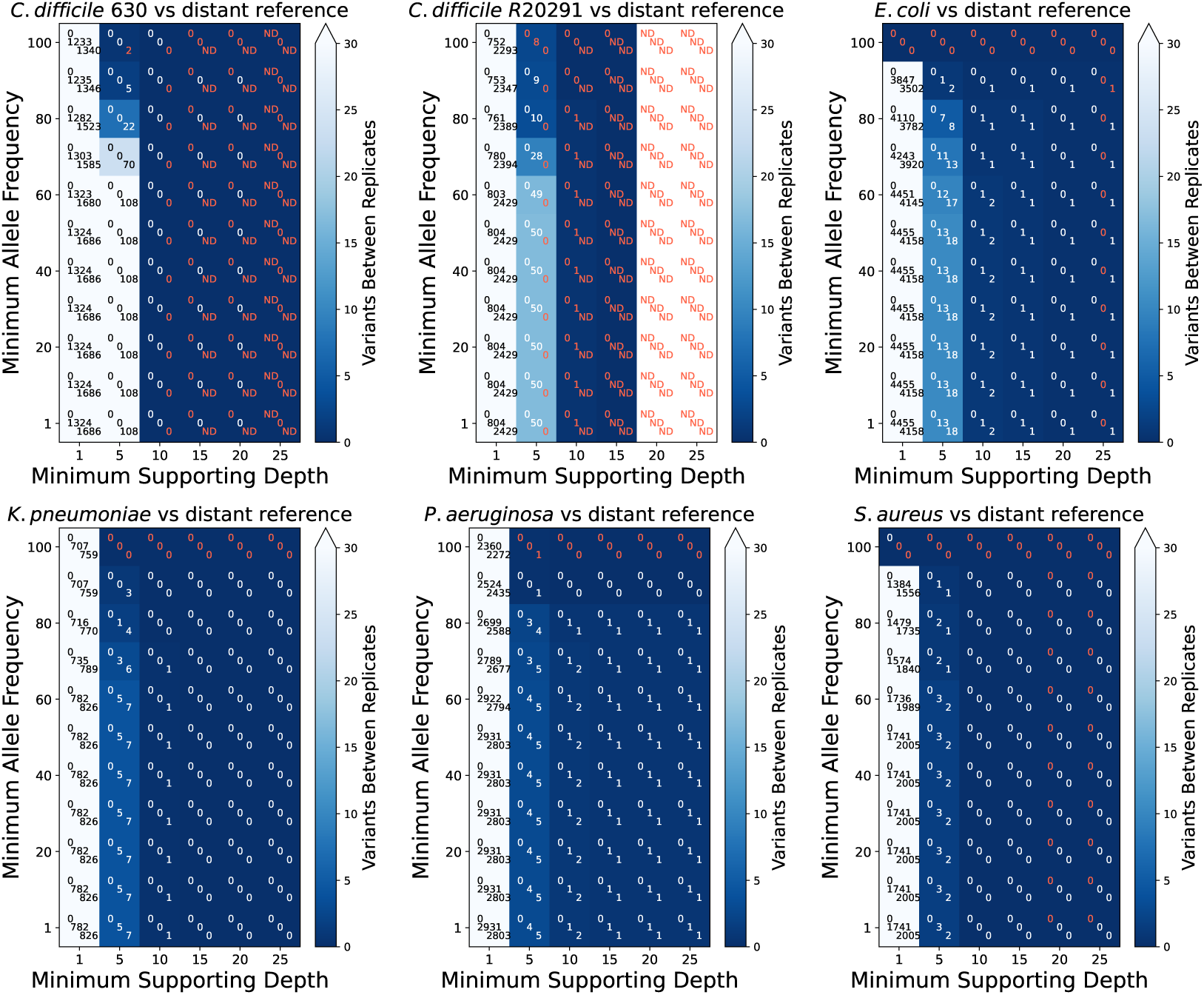
Number of variants between replicates of the reference set, when mapped to distant reference, by minimum frequency and read depth supporting each call. The number of variants for each replicate is shown in the centre of each cell and cells are coloured based on the number of variants found with darker tones representing fewer variants (theoretical minimum 0). Red text shows number of variants found in combinations where less than 75% of the reference genome was called, due to the loss of breadth of coverage resulting from increasing the supporting depth (Figure S9). This suggests that the fewer variants detected is likely caused by loss of data rather than optimal combination of variant frequency and supporting depths. Samples with less than the required depth are indicated by ND (no data). Figure S10 shows the same information for samples with higher sequencing depths.

Investigating the false positive variants when mapping the reads to the true reference in more detail (Table 2), most were in coding regions. In two cases (*C. difficile* 630 and *K. pneumoniae*) the same false positives were found in one of the replicates, but some were not reported due to low observed frequencies as per the ‘*OxBreaker*’ constraints, thus leading `snp`-`dists` to show different numbers of false positives when comparing the samples to the true reference (Figure 4). In most cases, the unique mutations occurred next to homopolymers (Table 2), context that `Dorado` has difficulty handling^26–28^, however the fact that false positives were present in all replicates, accumulation of mutations in the cultured strains cannot be excluded. There was a strong bias towards transversions in the false positive variants, particularly `C`→`A`. `C`→`G`, G→C, and `T`→`G` mutations (Figures S8A and B). This is consistent with prior findings^42^ and likely driven by the sequencing of methylated bacterial DNA^42,43^.

Overall, based on the self-reported read accuracy, we found between 0.55 and 3.5 errors per 100Kb in sequences with more than 75% coverage, with up to 400% variation in error estimates between replicates within species (Figure S8B). This error is consistent with prior estimates comparing ONT data sequenced with R10.4.1 chemistry against gold-standard Illumina datasets^43^. While inconsistencies between species are a known problem in ONT datasets^42^, the small number of false positives in the replicate reference set is therefore plausibly due to base-calling errors or accumulation of mutations with respect to the reference genome.

### Validating ‘*OxBreaker*’ with known clonal- and plasmid-driven outbreaks

To validate our pipeline, we re-analysed confirmed clonal^44^ and plasmid^45^ outbreaks from the literature. A clonal outbreak^44^, caused by *Staphylococcus capitis*, was sequenced with ONT chemistry R.9.4.1 and contained 40 clinical samples plus environmental strains isolated from neonatal intensive care units (NICU) in Oxfordshire (UK). The aim of the original study was to evaluate the contribution of patient-to-patient or environmental transmission to the cases of *S. capitis* bacteremias and carriage in infants admitted to NICU. The dataset also included 2 *S. capitis* strains sequenced as controls, *S. capitis* DSM 6717 and NCTC11045.

When first running the pipeline with the auto-discovery option, ‘*OxBreaker*’ found that 91.68% of the reads were identified as *Staphylococcus capitis* and, of these, 74.42% as *S. capitis* subspecies *capitis*—the only entry for this species in the `Kraken2` Microbial database. However, ‘*OxBreaker*’ halted the run warning that none of the accessible *S. capitis capitis* reference genomes were resolved, with all containing more than one contig. After feeding ‘*OxBreaker*’ *S. capitis* NCTC11045 as the reference genome, it was able to reproduce the phylogeny published in [44] (Figure 6A) with four main clades: 1) One at the root of the phylogeny containing NCTC11045, 2) a second clade containing the second reference used (DSM 6717), 3) a small clade containing environmental sample 2 and clinical sample 16, and 4) the biggest clade containing the remainder of samples—all of which were mostly closely related NRCS-A from `Kraken2`. ‘*OxBreaker*’ excluded 7 samples (environmental sample 5, and clinical samples 15, 22, 28, 33, 34 and 40) which did not have >75% of the genome mapped to the reference at minimum 10X depth.

**Figure 6.**
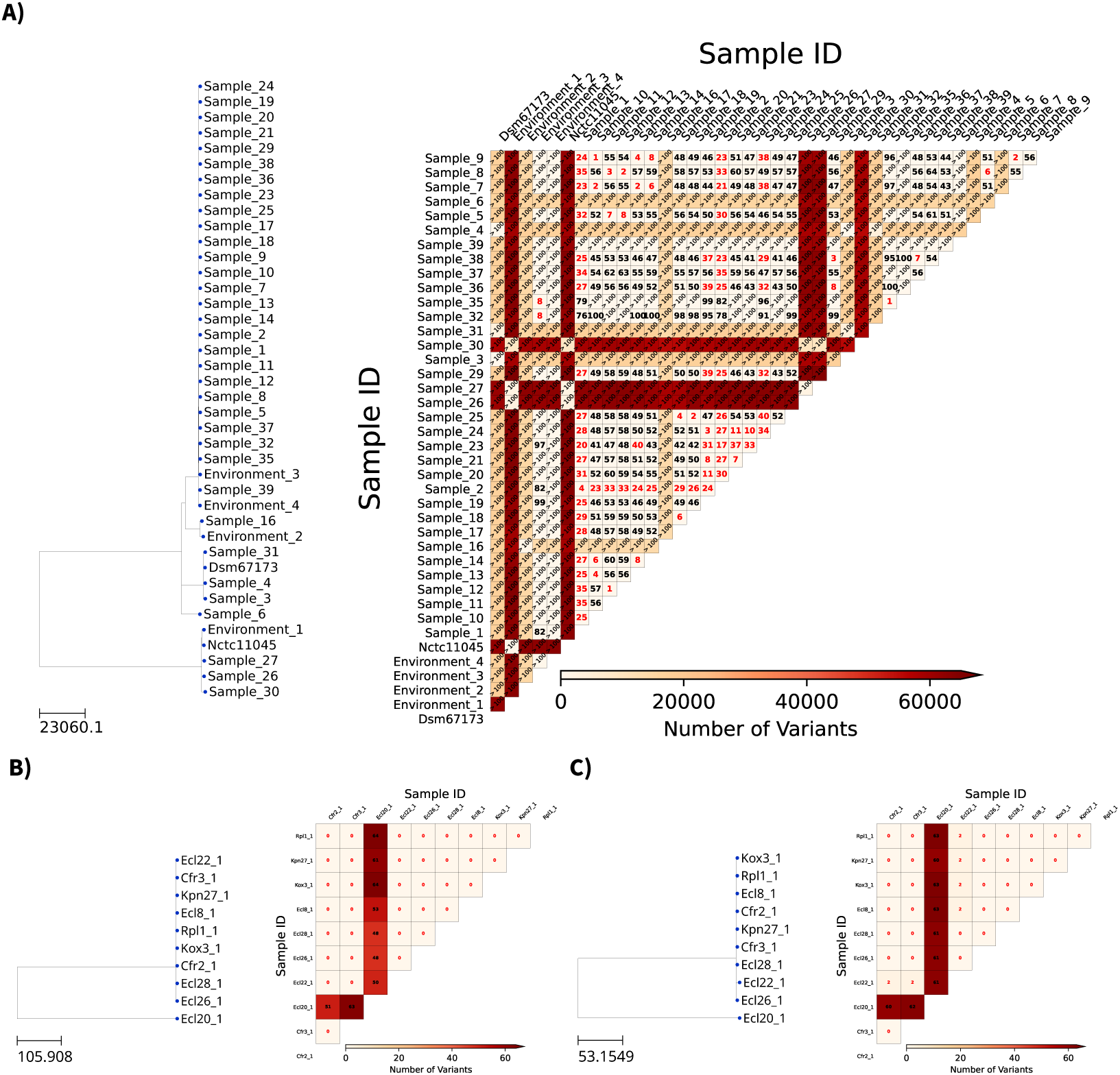
‘*OxBreaker*’ outputs for verified clonal *S. capitis* (A) and plasmid pQEB1/pQEB1_inv1 (B and C) outbreak datasets sequenced with R9.4.1 and R10.4.1 chemistries, respectively. For a given dataset, ‘*OxBreaker*’ produces a phylogeny (left), where the scale represents number of SNPs, and a SNP distance matrix (right) between the samples. For the distance matrix, darker red tones represent longer SNP distances between isolates, and lighter tones those more closely related. Red text indicates where there are fewer than 40 SNPs between two samples: For these, a table is produced showing the genomic context of the variants akin to Table 2. Black text indicates where there are more than 40 SNPs, but less than 100.

The accompanying distance matrix from ‘*OxBreaker*’ was also consistent with the report by Lees *et al.*^44^.‘*OxBreaker*’ identifies that many NICU cases of *S. capitis* are strains of a closely related cluster, but identifies that distances between isolates are not consistent with a simple point-source outbreak. Of note, two samples highlighted by Lees *et al.,* clinical samples 32 and 35, came from the same patient and are closely related to environmental sample 3 (door handle). The authors report a difference of 11 and 13 SNPs with respect to the environmental sample, respectively, and ‘*OxBreaker*’ reports 8 for both. SNP resolution with ‘*OxBreaker*’ is lower than that seen by Lees *et al.*^44^, which used a bespoke outbreak-specific reference genome. It is known that genomic diversity affects the accuracy of SNP detection in bacteria^46^. Overall, ‘*OxBreaker*’ pairwise distance estimates are strongly correlated with Lees *et al.*^44^ (Figure 7). In some comparisons, ‘*OxBreaker*’ estimated distances <20 SNPs where Lees *et al.* estimated ∼100 SNPs, for example between Samples 5 and 12 (8 SNPs), and 5 and 8 (6 SNPs). When the outbreak-specific, bespoke reference used by theses authors was provided to ‘*OxBreaker*’, these pairs showed distance estimates of 85 and 62 SNPs, respectively, more consistent with the earlier report (Figure 7B).

**Figure 7.**
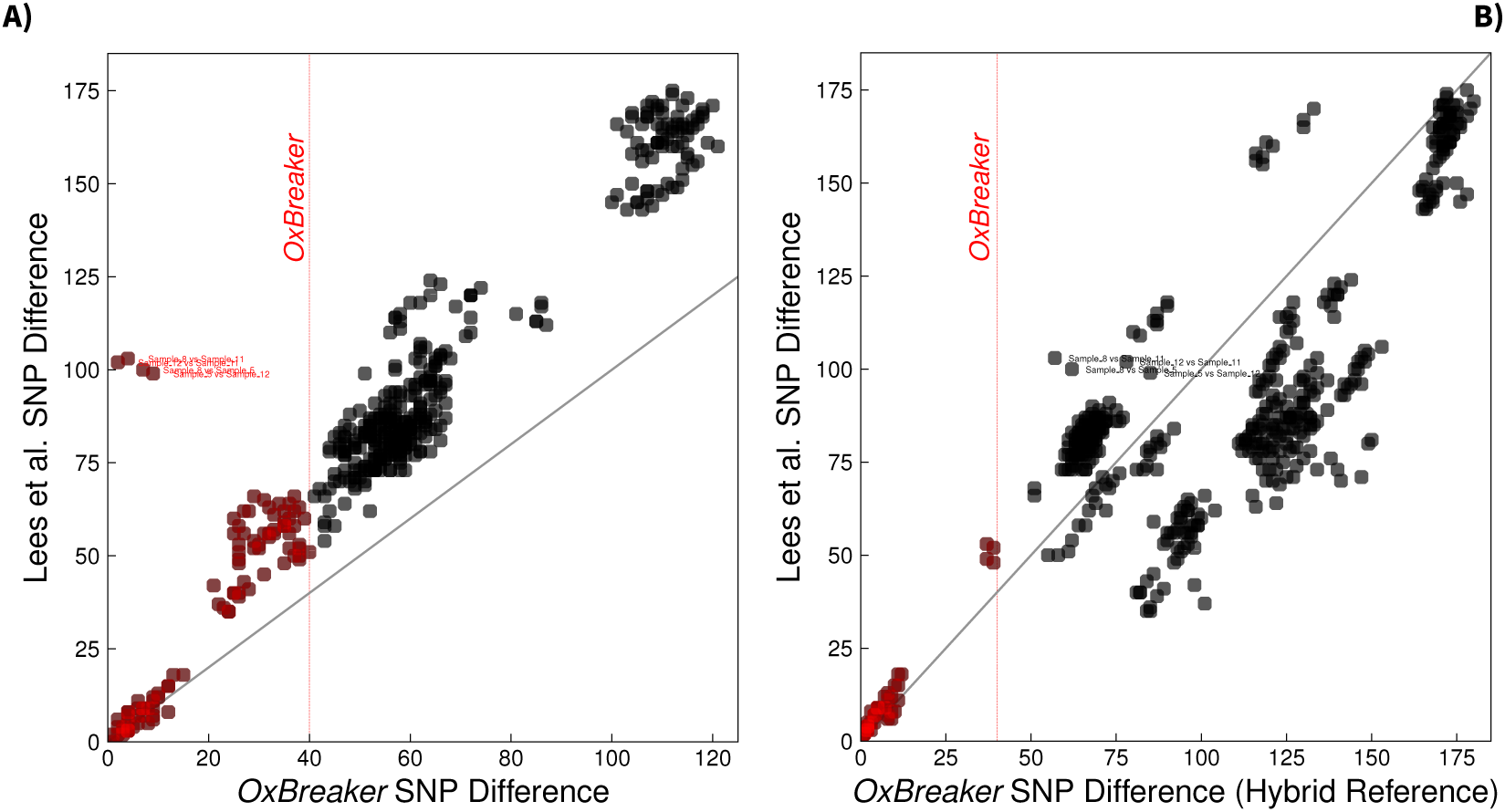
Comparison between the SNP distances reported by ‘*OxBreaker*’ and Lees *et al.*^44^ (A) and by ‘*OxBreaker*’ using the hybrid reference genome from [44] (B). Each dot represents a pair of samples analysed in both studies, with the SNP distance reported by ‘*OxBreaker*’ shown on the *x*-axis and the SNP distance by Lees *et al.* on the *y*-axis. The diagonal grey line represents the case where the distance in both cases is identical. Thus, a distance that is above this line means that more SNPs between isolate pairs were identified by Lees *et al.* whereas if they are below the line more SNPs between isolate pairs were identified by ‘*OxBreaker*’. Pairs of samples within a distance of 40 SNPs according to ‘*OxBreaker*’ are highlighted in red, with the red dashed lines noting this threshold. Pairs showing abnormal discordance between both methods, with ‘*OxBreaker*’ auto-discovered reference, are specifically labelled.

A plasmid outbreak^45^ was detected during the genomic surveillance of carbapenemase-producing Enterobacteriaceae at Birmingham’s Queen Elizabeth Hospital. The KPC-associated plasmid was first found in an *Enterobacter cloacae* (Ecl) isolate, prompting further sequencing with 11 clinical samples isolated from 10 patients, and sequenced using ONT with R10.4.1 chemistry. Representative bacterial hosts included three sequence types of *Citrobacter freundii* (Cfr), two of *Enterobacter cloacae*, and three species of *Klebsiella*: *K. pneumoniae* (Kpn), *K. oxytoca* (Kox), and *K. planticola* (Kpl) isolated in 7 different hospital wards. For our analysis, we fed ‘*OxBreaker*’ the two plasmid references provided by the authors, pQEB1 (Figure 6B) and pQEB1_inv1 (Figure 6C). For both references, ‘*OxBreaker*’ identified the original sample prompting the genomic surveillance study (Ecl-20) and the distance between all samples except Ecl-20 were between 0–2 SNPs, depending on the reference plasmid used. While a more nuanced analysis would be required to pinpoint the plasmid that sparked the outbreak, ‘*OxBreaker*’ provides an output that would inform urgent action for infection prevention and control.

### Application to potential outbreaks caused by *C. difficile*, *Streptococcus gallolyticus*, and *Staphylococcusaureus* datasets

Following this validation of ‘*OxBreaker*’ using examples from the literature, we moved on to evaluate previously unpublished sequences from potential outbreaks of *C. difficile*, *Streptococcus gallolyticus*, and *Staphylococcus aureus* in the Oxford University Hospitals NHS Foundation Trust, Oxford, UK.

The first contains 30 clinical *C. difficile* isolates from stool samples of patients admitted to hospital, with 6 controls added to the sequencing run comprising 3 replicates of each *C. difficile* 630 and *C. difficile* R20291 for a total of 36 samples analysed. The outbreak investigation was motivated by concern about increasing case numbers in the trust. Using the reference auto-discovery, the closest reference was *C. difficile* 630. with Kraken2 identifying 96.0% of the reads as *C. difficile* and 2.88% as Enterobacterales. Of those identified as *C. difficile*, the strain with most reads (2.5%) was ATCC 43255. However, its genome remains unresolved, having 34 scaffolds and therefore being unsuitable for phylogenetic analysis. After iterating through other potential *C. difficile* reference strains, the pipeline selected *C. difficile* 630 as the first one with a resolved genome. Figure 8A shows that the *C. difficile* 630 replicates (samples 01, 49, one technical and one biological replicate), and R20291 biological replicates 02, 50 were 0 SNP different from each other, as expected (*C. difficile* 630 and R20291 technical replicate 65 and 66 excluded by the pipeline due to low sequencing depth). The next most closely related pair of isolates (samples 52 and 53) were 43 SNPs apart, with the remaining pairs being all >100 SNPs. Hence, the samples did not show evidence of patient-to-patient transmission or a common source. However, 21 clinical samples and two references were excluded from this analysis because they did not have >75% of the genome mapped to the reference at 10X depth—consistent with poor sequencing yields.

**Figure 8.**
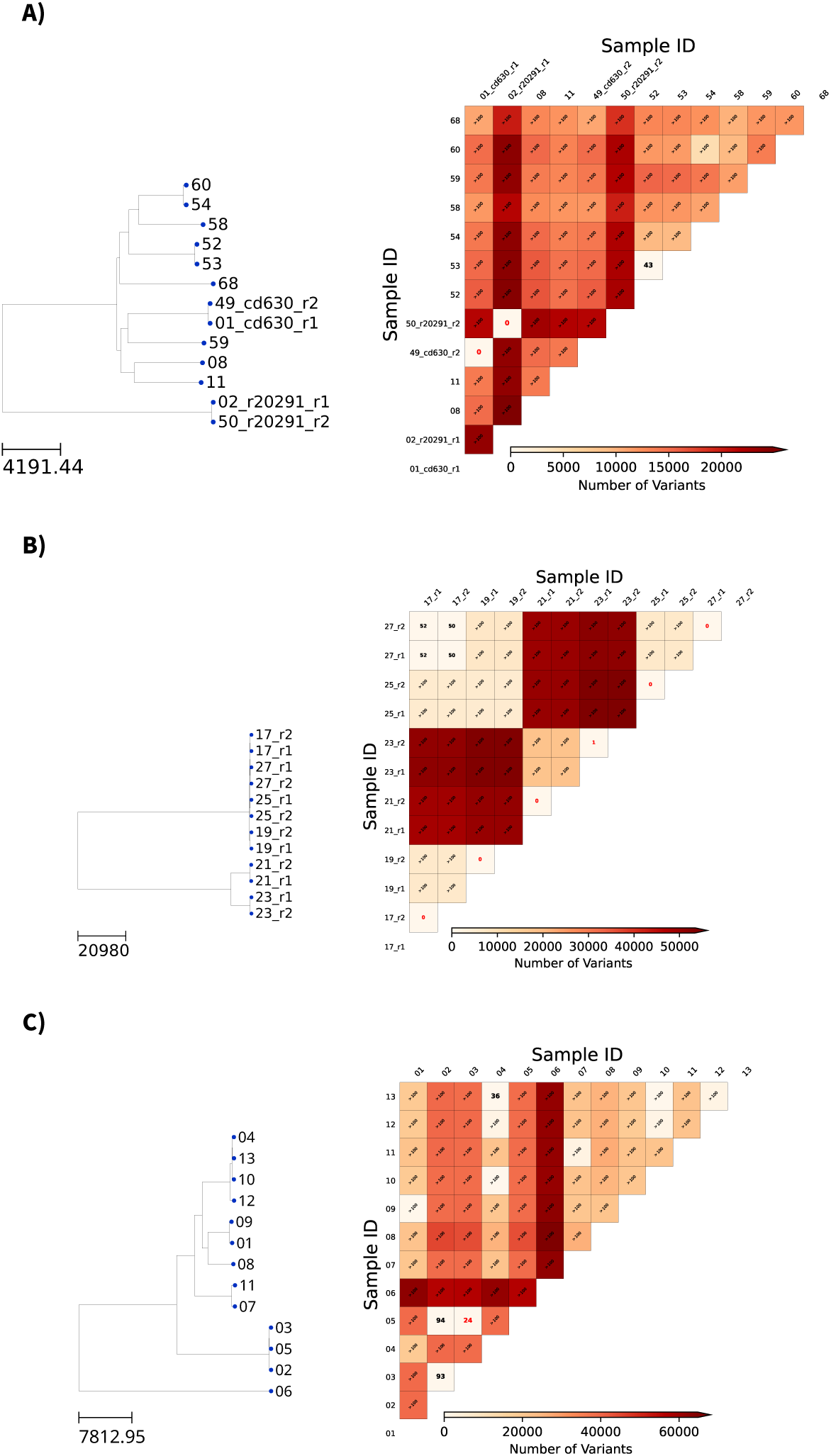
‘*OxBreaker*’ outputs for suspected *C. difficile* (A), *S. gallolyticus* (B), and *S. aureus* outbreak datasets sequenced with R10.4.1 chemistry. For a given dataset, ‘*OxBreaker*’ produces a phylogeny (left), where the scale represents number of SNPs, and a SNP distance matrix (right) between the samples. For the distance matrix, darker red tones represent longer SNP distances between isolates, and lighter tones those more closely related. Red text indicates where there are fewer than 40 SNPs between two samples: For these, a table is produced showing the genomic context of the variants akin to Table 2. Black text indicates where there are more than 40 SNPs but less than 100.

The second potential outbreak contains 6 clinical *S. gallolyticus* isolates from patients with endocarditis after transcatheter aortic valve implantation. The possibility of a common source for these infections was raised due to unexpected prevalence of endocarditis cases with this organism, which is typically associated to health issues in rumiants^47^. Isolates from blood cultures, between December 2022 to March 2024, were sequenced using ONT (see methods) and each isolate had one technical replicate. As Figure 8B illustrates, ‘*OxBreaker*’ identified two main clades in this dataset: One containing samples 21 and 23, and a bigger clades with the remainder of the isolates. While only the replicates are within 0–1 SNPs, overall the dataset gives evidence against a common source for the infections with >100 SNPs between most pairs. However, ‘*OxBreaker*’ identified a 50–52 SNPs difference between samples 17 and 27, reflecting the ability of the pipeline to resolve close relationships between isolates.

The last potential outbreak used to test ‘*OxBreaker*’, was an investigation conducted after observing multiple cases of methicillin-resistant *S. aureus* (MRSA) within one clinical unit in the Trust. With all cases showing phenotypic resistance to gentamicin, a trait only seen in 25% of local MRSA isolates, the IPC teams questioned whether a newly dominant lineage was emerging. We sequenced 2 MRSA isolates from bloodstream infection taken between between May and September 2025; 10 gentamicin-resistant MRSA isolates, also from bloodstream infections, taken between September 2022 and September 2025; along with a reference strain (MRSA-252). As Figure 8C shows, the cases showed significant diversity suggestive of multiple circulating lineages. However, ‘*OxBreaker*’ identified two clusters within these samples: One containing isolates 4 and 13, 36 SNPs apart, and another containing isolates 3 and 5, 24 SNPs apart. These were within the 20–40 SNP thresholds proposed to exclude transmission in MRSA^37,38^. Following an epidemiological review, we found only one of these isolates came from the clinical unit of concern, and neither pair had shared ward space or clinical teams. Thus, in this case, ‘*OxBreaker*’ was able to rapidly generate evidence against the hypothesis of a point-source outbreak, and allowed a more focused epidemiological review by narrowing IPC action to two possible pairs of cases.

## III. DISCUSSION

ONT sequencing is an attractive option for informing IPC practice given its potential for rapid sequencing, increasing application in clinical settings given the improved accuracy of ts R10.4.1 chemistry—particularly with its super-accuracy (SUP) base-calling algorithm—and low overheads. However, sequence analysis requires specialised expertise, creating a bottleneck for the implementation of ONT sequencing as a routine tool available to practitioners. Here we developed and evaluated a species-agnostic pipeline, ‘*OxBreaker*’, which, through its graphical user interface, could be used to fill this gap in the adoption of ONT sequencing for outbreak investigation. Through a systematic screening of parameters relating to mapping quality, read depth and variant frequency, and use of a set of biological and technical replicate sequences from reference strains of 6 HCAI pathogens, we found a combination that is accurate, as measured by minimal false positives in the replicate reference set, and yet applicable to multiple species and even plasmids, regardless of the chemistry used if the SUP base-calling algorithm is used. In other words, ONT sequencing could be suitable for tracking outbreaks and informing counter-measures for IPC interventions when combined with ‘*OxBreaker*’.

Perhaps the most interesting finding is the lack of improvement, in terms of reducing false positives, of requiring increased sequencing depth above 20X to report a variant. The ongoing challenge with reproducible ONT sequencing yield caused the variability in sequencing depths in *K. peumoniae* or *S. aureus* MRSA252. This, in turns, offers an opportunity to assess what outputs from real-world sequencing datasets may look like, and highlights the stability of the false positives detected by ‘*OxBreaker*’. This shows both the robustness of our pipeline to variations in sequencing depth and mitigates the need for resequencing, which would allow the multiplexing of samples to further reduce costs and turnaround times.

However, there are some limitations. First, is the impact of lower than expected sequencing yields from GridION flowcells, as Figure S8B illustrates. While we mitigate this limitation by optimising the parameters of ‘*OxBreaker*’ to allow the analysis of samples with low depth (at least 10X), runs may still fail without prior warning and specific samples may be excluded from analyses—limiting inference for IPC actions. Such was the case for two samples from the replicate reference set, corresponding to one of the replicates of each *C. difficile* 630 and *C. difficile* R20291. And, perhaps more importantly, the exclusion of 21/30 samples (70%) from the potential *C. difficile* outbreak dataset. This was caused by flow cells with lower yields than expected following an internal investigation. A second limitation, with mutation bias toward `C`→`G` and *C*→*G* mutations compared to the reference, lies with methylated genomes. Prior research^48^ has shown that such mutations are common signature of methylated nucleotides, that are miss-called by `Dorado`^26–28^. Using a version of `Dorado` trained to detect methylated nucleotides could help further reduce the number of false positives with respect to the reference. But these models cannot be run concurrently since each model is specific to certain genomic modifications. However, given that the false positives we found compared to the reference were consistent across biological and technical replicates, it is also possible that our cultured strains have accumulated these mutations during laboratory passage over the years.

A third limitation is the availability of fully resolved assemblies that can be used as a reference by ‘*OxBreaker*’ through its auto-discovery mode, which depends on microbial classification databases. Alternatively, if there are multiple entries for a given species, then ‘*OxBreaker*’ may select a sub-optimal reference. For example, in the *C. difficile* potential outbreak, most reads were identified as *C. difficile* ATCC 43255 but ‘*OxBreaker*’ selected strain 630. This is because it was the first `Kraken2` entry with only one contig whereas ATCC43255 has 23 contigs. To investigate the risk of errors arising from the specific choice of reference, we validated ‘*OxBreaker*’ using both the true and distant references. This confirmed no increase in the number of false positives between replicates—since the reference is only used to aid assembly of the reads, not in the analysis—but demonstrated that lower resolution is achieved with a more distant reference where sequencing depth is marginal. Nevertheless, the size of the Kraken2 microbial database (30GB) is a potential limitation for running the auto-discovery routine on local IT infrastructure in some hospitals. This limitation is mitigated by the ability to provide a known reference supported by laboratory results—albeit this requires some knowledge of how to find and download genome sequences.

Although designed for use with outbreaks of clonal HCAI, we demonstrate that ‘*OxBreaker*’ can also be used to identify plasmid-driven outbreaks where evolution is predominantly driven by mutation. In contrast to auto-discovery of chromosomal references, however, auto-discovery of plasmid references is a much larger challenge: There is no `Kraken2` equivalent for plasmids. Combined with their sheer number and variety, the identification of plasmid-mediated outbreaks could be even more complex. For example, in the case of pQEB1 and pQEB_inv1, Moran *et al.*^45^ established that pQEB_inv1 derives from an IS26 inversion event of pQEB1; but the sequences are otherwise almost identical. ‘*OxBreaker*’ was not designed to use with plasmids, and so is not able to provide this resolution depth. However, what it did do, as shown in Figures 6B and C, is to confirm the existence of a plasmid-mediated outbreak. For precise plasmid identification in this case, for example, it could be implemented in ‘*OxBreaker*’ the *de novo* assembly of the reads as an alternative in a future release, given the reduced genome sizes involved. While a database would still be required for rapid plasmid identification, this approach would generate a plasmid genome that can be used to study its origin further.

A key question in applying pathogen sequencing to outbreak detection is considering the number of SNP differences that can be considered consistent with transmission in an outbreak. Based on previous research, these vary by organism, sequencing technology and analytic approaches^38,49–51^. The upper end of this range encompasses the lower bounds of some species specific thresholds, lending some uncertainty to the ability of ‘*OxBreaker*’ to exclude direct transmission in marginal cases. Whilst there is interest in identifying direct patient-to-patient transmission, sampling for outbreak investigations will likely be incomplete since samples are often taken for clinical purposes, rather than routinely screened. There is also a lack of routine, environmental sampling. ‘*OxBreaker*’ can help IPC teams identify whether a cluster of cases has close genetic relationship, supporting epidemiological investigation—particularly in the early stages of an outbreak where such link may be more difficult to establish. Moreover, the ability to exclude transmission or a common source of infection may be as important as supporting a potential outbreak, as it helps focus the available resources on exploring other contributors to the infections. In the case of our *C. difficile* investigation, for example, to focus on antimicrobial stewardship.

The literature on real-time ONT sequencing for outbreak detection relatively sparse^52–54^, all with pipelines that are difficult to scale given their requirement for high sequencing depths and difficult to implement given the lack of accessibility to the pipelines. For example, the code by Vereecke *et al.*^53^ is not accessible, despite using open-source tools, and requires extensive command-line expertise. Similarly, the pipeline by Cottingham *et al.* requires command-line expertise, is not available to download, and requires a sequencing depth of at least 60X to confidently call variants. SKA2^52^ is a promising tool for outbreak analysis, but requires extensive command-line expertise. The platform from SOLU Genomics^20^ is the only tool we found that offers an intuitive web-based interface for outbreak analysis, however, it incurs further costs per sample—additional to the sequencing costs—and requires uploading the samples to external servers and thus requires continuous internet connection. This step adds a layer of complexity since hospital personnel must ensure compliance with local laws and IT outages can prevent access to the data, a known pitfall of internet-based solutions in healthcare^55,56^. The costs of short-read, whole genome sequencing can impose a substantial financial burden on local hospitals and thus can be expensive to implement widely. However, in few instances where genomic surveillance was thoroughly implemented for outbreak analysis^3,57^ relied on bespoke pipelines— like (EDS-HAT)^3^—that combined short-read sequencing with hospital-level data on ward movements and machine learning.

We circumvent these issues with ‘*OxBreaker*’ by enabling genomic surveillance using ONT sequencing. This technology shows promise to facilitate routinely sampling for sequencing. ‘*OxBreaker*’ provides actionable information for infection prevention and control teams—including the reallocation of resources—at no extra cost, relies on open-access tools, is accessible to both developers and end-users through its GUI, can be used off-line, while retaining data ownership. Despite our promising results, particularly where ‘*OxBreaker*’ could replicate a *S. capitis* study for IPC purposes as illustrated in Figure 7, further studies are still needed to demonstrate the clinical and health economic impacts of implementing ‘*OxBreaker*’ alongside ONT sequencing for IPC.

## IV. METHODS

### Sample sequencing

For six reference strains, DNA was extracted from frozen laboratory stocks. Bacteria were stored at −80*°*C in nutrient broth plus 10% glycerol. These were cultured on Colombia Blood Agar (CBA) at 37*°*C overnight, and single colony was sub-cultured overnight on CBA at 37*°*C. For outbreak investigations of *S. gallolyticus* and *S. aureus* from blood culture, isolates were retrieved from storage in glycerol stocks at −80*°*C and cultured as for the reference strains. Clinical *C. difficile* isolates were cultured from stool identified for investigation, with a loop of stool vortexed in 70% ethanol and incubated at room temperature for 15 minutes: 5of that mix was cultured on Cefoxitin Cycloserine Egg Yalk (CCEY) agar anaerobically for 3–7 days until growth was seen. From this culture, approximately 6 colonies were sub-cultured anaerobically at 37*°*C for 48h. For all cultured isolates, DNA was extracted using the QIAGEN Genomic Tip 100G-1 kit and for the clinical *C. difficile* samples we used QuickGene DNA Tissue Extraction (ADS Biotec) in accordance with the manufactures’ instructions. DNA quality was assessed with the Qubit Flurometer and TapeStation after extraction.

All sequencing of our reference and potential outbreak datasets was performed on an ONT GridION device using a 5kHz sampling rate and R10.4.1 flow cells. Libraries were prepared using the Rapid Barcoding Kit (RBK114.96) and run for 10h (clinical *C. difficile*) and 72h for all other strains. Two *C. difficile* reference strains were sequenced alongside clinical strains in multiplexed libraries of 12 extracts per flow cell. The remaining four references (*E. coli, K. pneumoniae, S. aureus, P. aeruginosa*) strains sequenced were multiplexed and run on a single flow cell. Additional reference strain sequences were generated previously in [43].

### Reference sequences

The accession code for the reference genomes used are NZ_CP010905 (*C. difficile* 630), NZ_CP029423 (*C. difficile* R20291), NZ_CP051263 (E. coli CFT073), CP000647 (*K. pneumoniae* MGH78578), NC_002516 (*P. aeruginosa* PA01), and BX571856 (*S. aureus* MRSA252) from the National Center for Biotechnology Information (NCBI) database.

### Pre-processing data

We used ONT’s `Dorado` version 1.0.2 for base-calling R10.4 the reads using super accuracy model version 5.2.0 (dna_r10.4.1_e8.2_400bps_sup@v5.2.0), with standard options and saving the output as FASTQ files (`--emit-fastq`). These files are piped to gzip to generate the final compressed FASTQ file. We ran this process in a workstation with an Intel Xeon Gold 5217 (16 threads), 192 GB of memory, and four NVIDIA GeForce RTX 2080 Ti running Ubuntu 22.04.5 LTS.

We ran ‘*OxBreaker*’ on a workstation equipped with an Intel Xeon W-2295 (36 cores) limited to 16 cores, and 128GB of memory. We monitored memory usage and ‘*OxBreaker*’ never surpassed 13GB in our datasets when using a known reference genome, and 32GB with auto-discovery when the 30GB is loaded into memory by `Kraken2` with option `--memory-mapping`, suggesting it could be run in equipment readily available in hospitals in the UK. However this compute requirement may vary as a function of sequencing depth as samples with deeper sequencing produce larger files, and larger databases may require more memory. Base-called read summary statistics were generated as part of the quality assessment run by ‘*OxBreaker*’, using NanoPlot^58^ version 1.44.1 with standard options.

GridION will produce multiple files per barcode as part of the sequencing process, resulting in multiple FASTQ from `dorado`. ‘*OxBreaker*’ can merge these FASTQ to produce one single FASTQ file per barcode if the filename convention entered in GridION has the barcode number first (i.e. `barcodeXX_runIdentifier_hash.fastq.gz`, where XX corresponds to the barcode number). Following this, the resulting `barcodeXX.fastq.gz` files are stored in the directory `reads_merged/`, to preserve the originals, and are used downstream by ‘*OxBreaker*’.

### Funding

This study was funded by the National Institute for Health Research (NIHR) Biomedical Research Centre, Oxford, and also supported by the NIHR Health Protection Research Unit in Healthcare Associated Infections and Antimicrobial Resistance (NIHR207397), a partnership between the UK Health Security Agency (UKHSA) and the University of Oxford. The views expressed are those of the authors and not necessarily those of the NIHR, UKHSA or the Department of Health and Social Care. ASW is an NIHR Senior Investigator.

### Availability of data and materials

Sequencing data corresponding to the replicate reference dataset, as well as potential outbreak datasets for *C. difficile*, *S. gallolyticus*, and *S. aureus* were deposited in the European Nucleotide Archive under project PRJEB108014. A basic NextFlow implementation of ‘*OxBreaker*’, licensed under the Lesser General Public License (LGPL) v2.1, can be found at https://github.com/oxfordmmm/OxBreaker.

### Ethics

The use of cultured isolates from clinical samples was approved by an NHS Research Ethics Committee (REC), reference 17/LO/1420. This approval covers the use of surplus materials from routinely collected samples processed by the Oxford University Hospitals NHS Foundation Trust Clinical Microbiology laboratory, and linkage of these samples to de-identified patient information, for research into the development and evaluation of sequencing tests without patient-level consent.

## Acknowledgements

The authors are grateful to Dr Hieu Thai (Nuffield Department of Medicine, University of Oxford) for the procurement of virtual machines for the calibration of ‘*OxBreaker*’.

## I. SUPPLEMENTARY FIGURES

**Figure S1.**
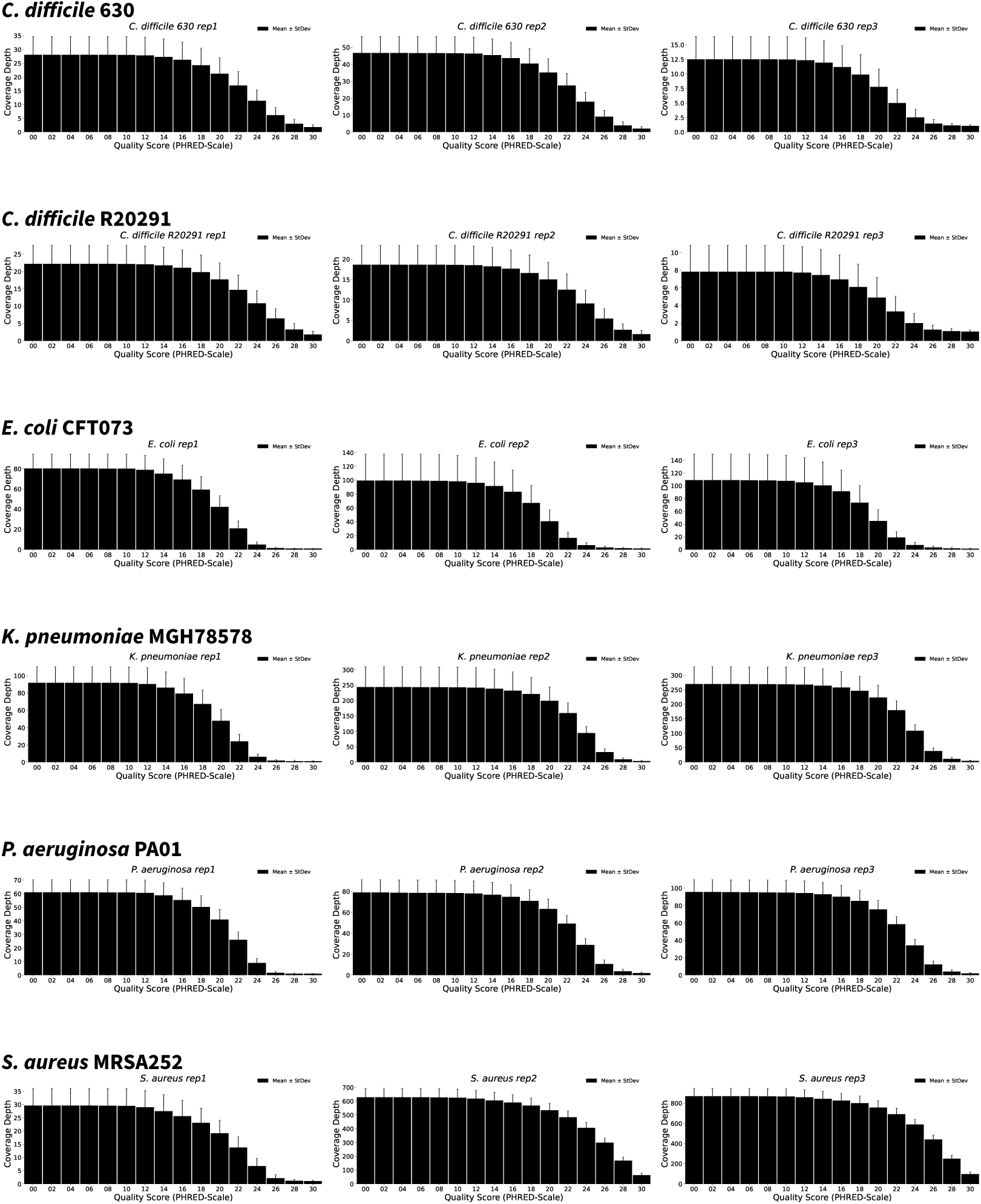
Raw read depth with increasingly stringent quality thresholds in the replicate reference set. For each sample in Table 1, we estimated the depth of coverage of the reference genome resulting from filtering reads prior to mapping them to the reference. Each figure shows the mean coverage ± standard deviation after filtering the reads between Phred quality scores 0–30.

**Figure S2.**
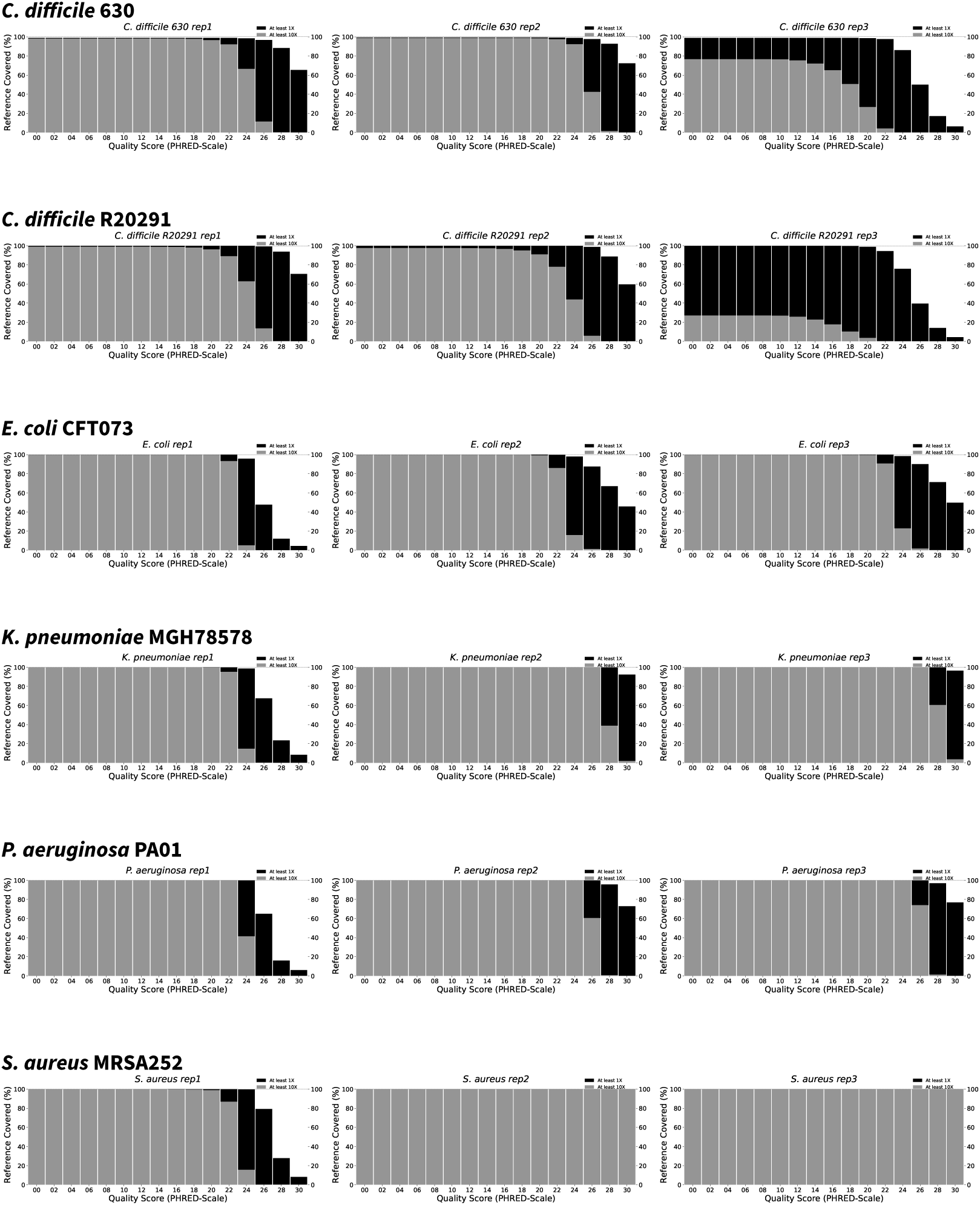
Raw coverage breadth with increasingly stringent quality thresholds in the replicate reference set. For each sample in Table 1, we estimated the breadth of coverage of the reference genome resulting from filtering reads prior to mapping them to the reference. Black columns note the percentage (*y*-axis) of the reference with a read depth of at least 1X, whereas the grey columns show the percentage of the reference with a read depth of at least 10X.

**Figure S3.**
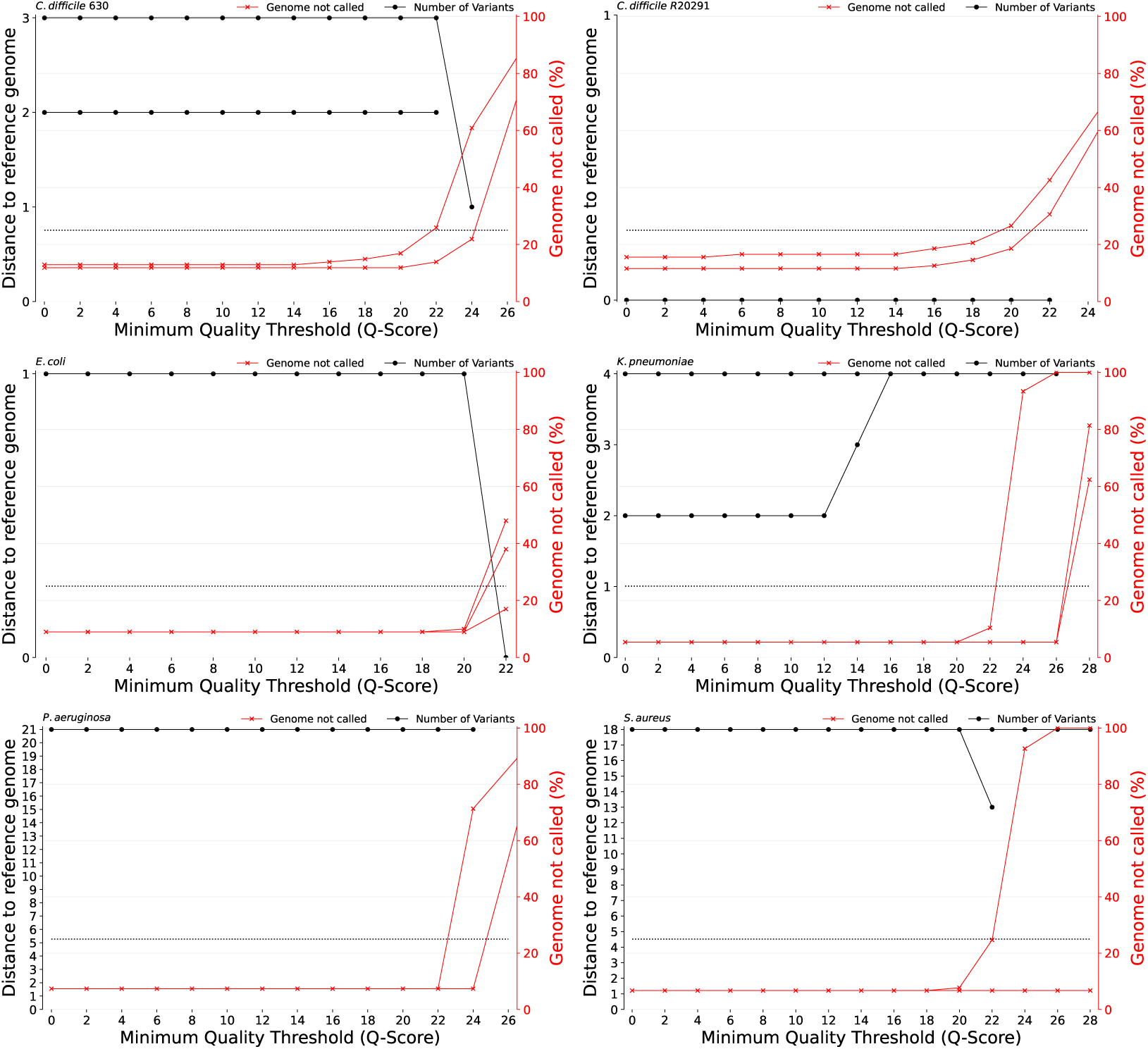
Number of variants detected and percentage of the reference genome not covered as a function of quality-score filtering thresholds in the replicate reference set. The variants found in each isolate after mapping to their true reference are shown in the *y*-axis at the left species after filtering reads below a quality score of 0–28 (with 0 meaning no filtering). In red, the *y*-axis on the right shows the percentage of the genome that was not covered as a consequence of this read quality-score filtering. The horizontal, dotted line in black represents the maximum masking allowed before ‘*OxBreaker*’ excludes the sample from downstream processing.

**Figure S4.**
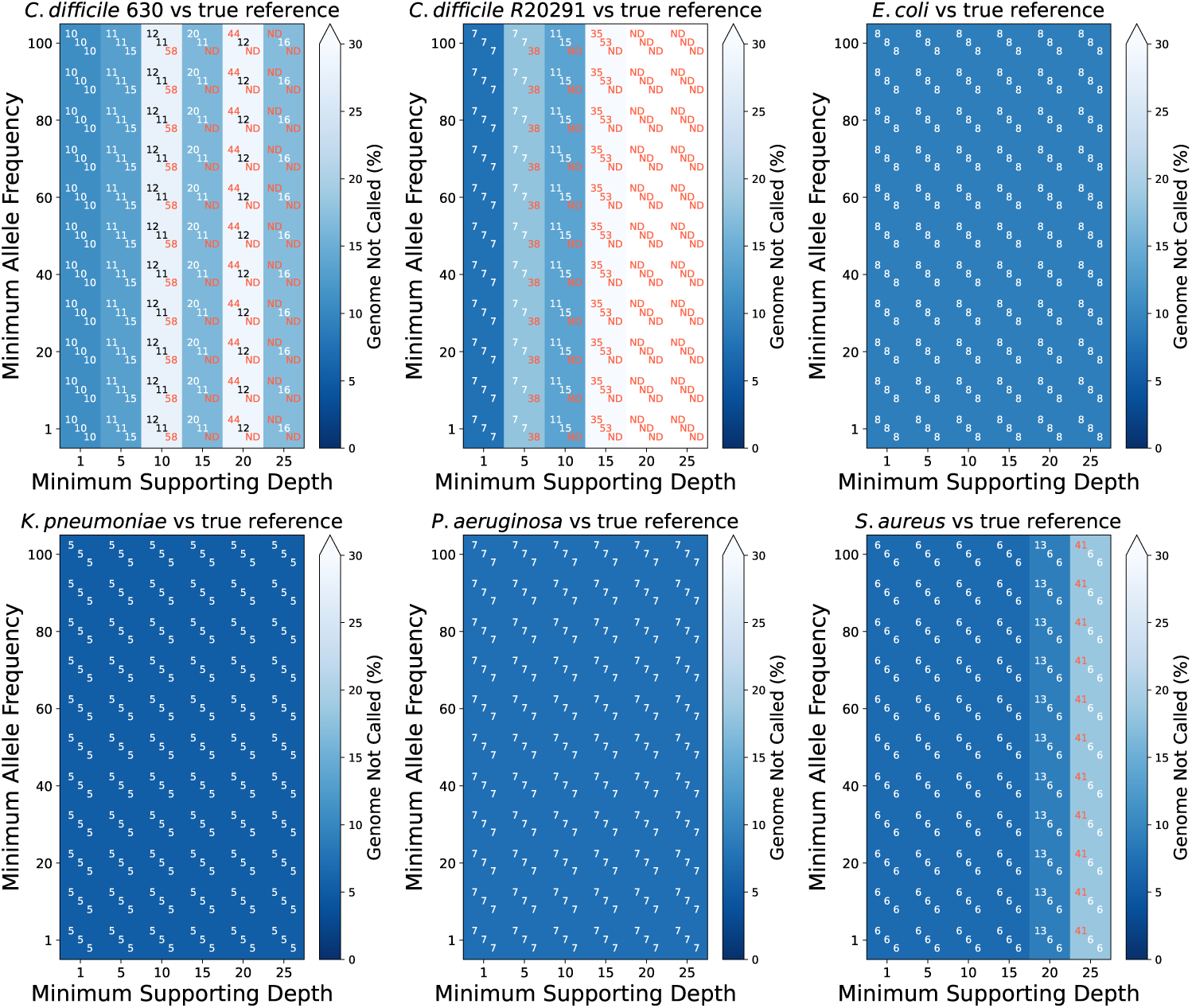
Percentage of genome not called in the replicate reference set when mapped against the true reference genome. The percentage of genome not called for each replicate is shown in the centre of each cell, and cells are coloured when the percentage is above 25%, those with most percentage of their genome available for analysis being represented with darker tones (theoretical minimum 0). Samples with less than the required depth are noted as ND (no data).

**Figure S5.**
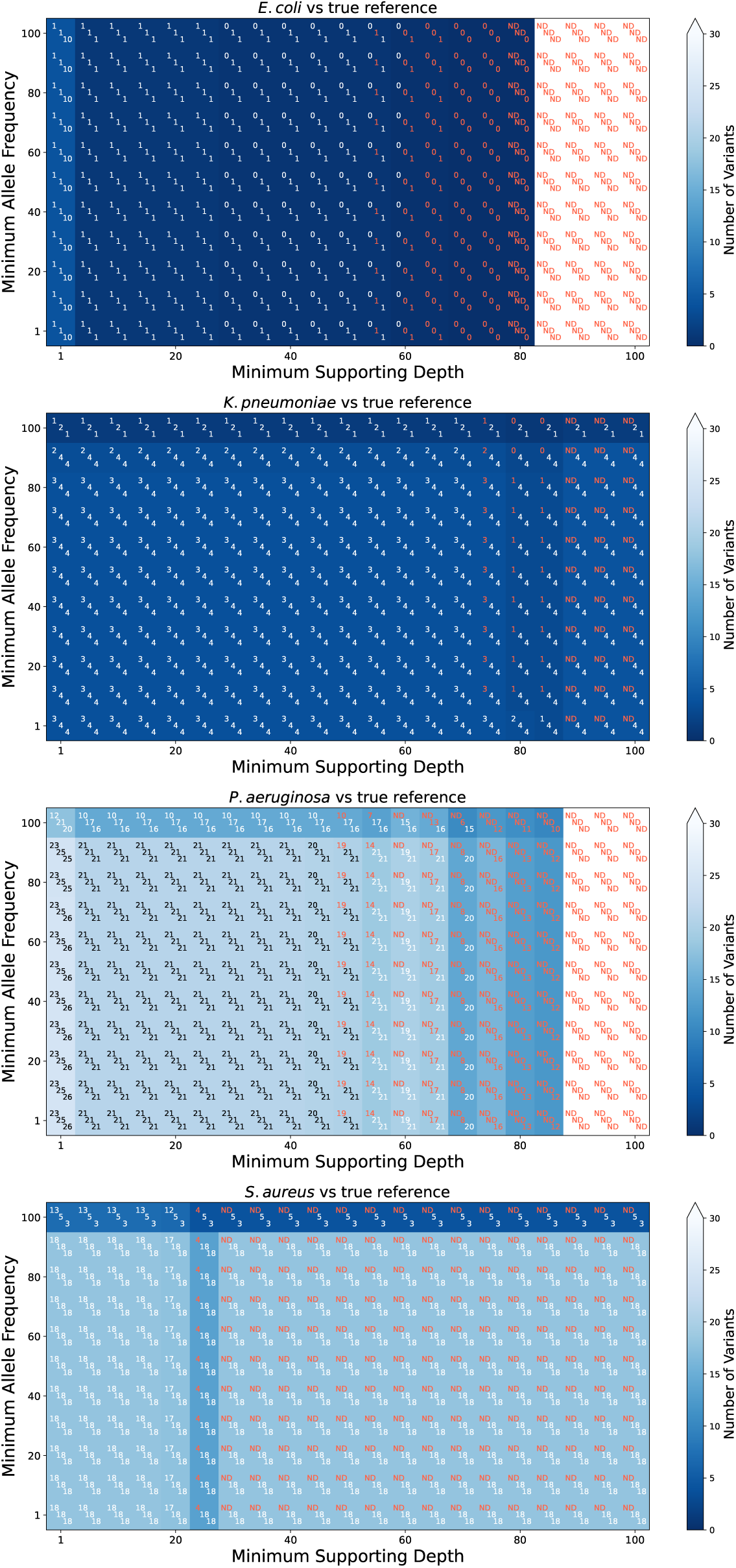
Number of false positives when mapping against the true reference in replicate reference set isolates with depth >50X optimise minimum frequency and minimum read depth. The number of variants for each replicate is shown in the centre of each cell, and cells are coloured based on the number of variants found with darker tones representing fewer variants (theoretical minimum 0). Due to the loss of breadth of coverage resulting from increasing the supporting depth (Figure S2), red text denotes the number of variants found in combinations where less than 75% of the reference genome was called. Samples without depth data available are noted as ND (no data). This serves as a means to warn that the fewer variants is likely caused by loss of data rather than optimal combination of variant frequency and supporting depths.

**Figure S6.**
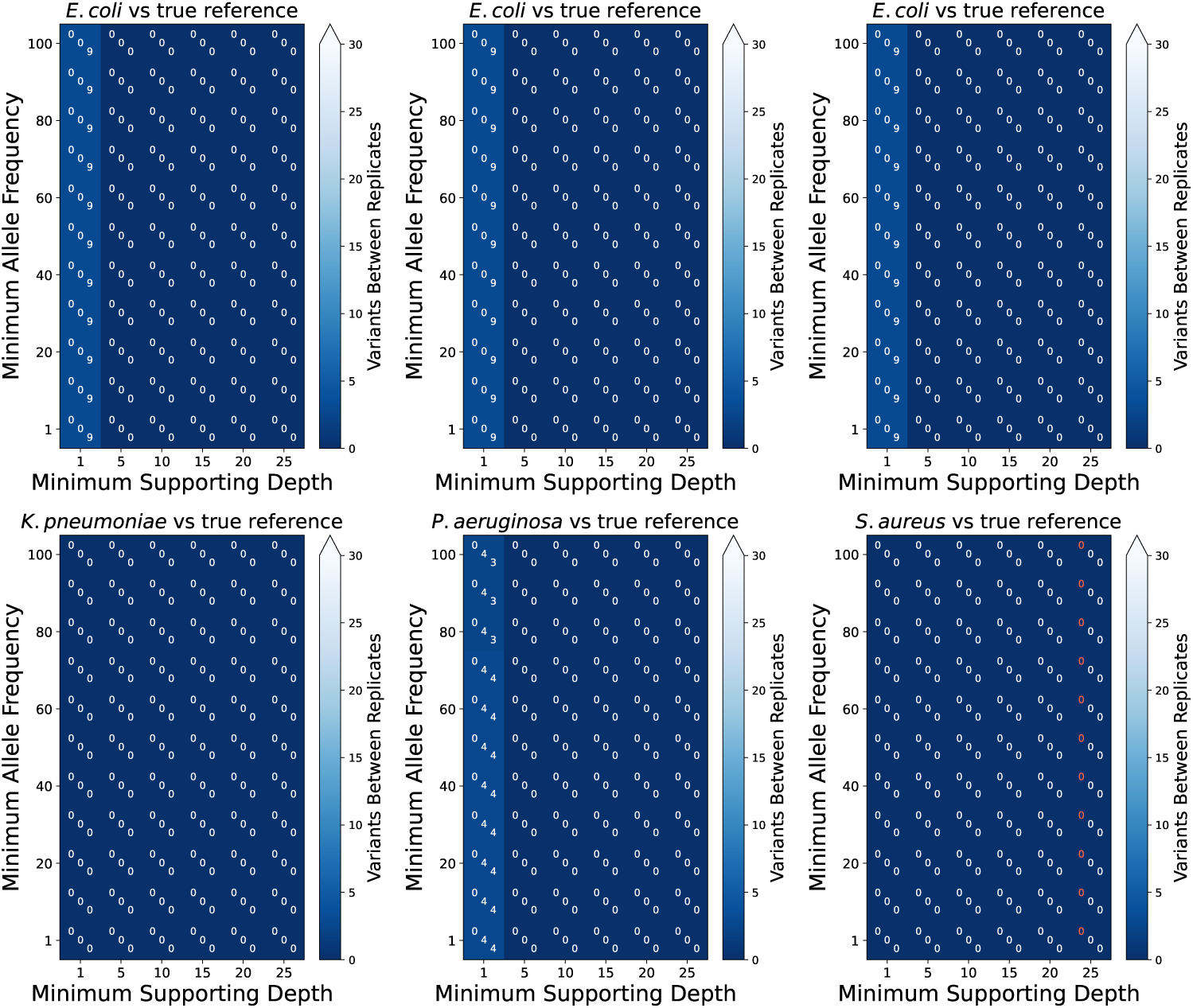
Number of variants between replicates of the replicate reference set, when mapped to the true reference, by minimum frequency and minimum read depth supporting each call. The number of variants for each replicate is shown in the centre of each cell, with the three numbers comparing replicate 1 *vs* 1, 1 *vs* 2, and 1 *vs* 3, and cells are coloured based on the number of variants found with darker tones representing fewer variants (theoretical minimum 0). Due to the loss of breadth of coverage resulting from increasing the supporting depth (Figure S4), red text denotes the number of variants found in combinations where less than 75% of the reference genome was called, suggesting that the fewer variants detected is likely caused by loss of data rather than optimal combination of variant frequency and supporting depths. Samples without depth data available are noted as ND (no data).

**Figure S7.**
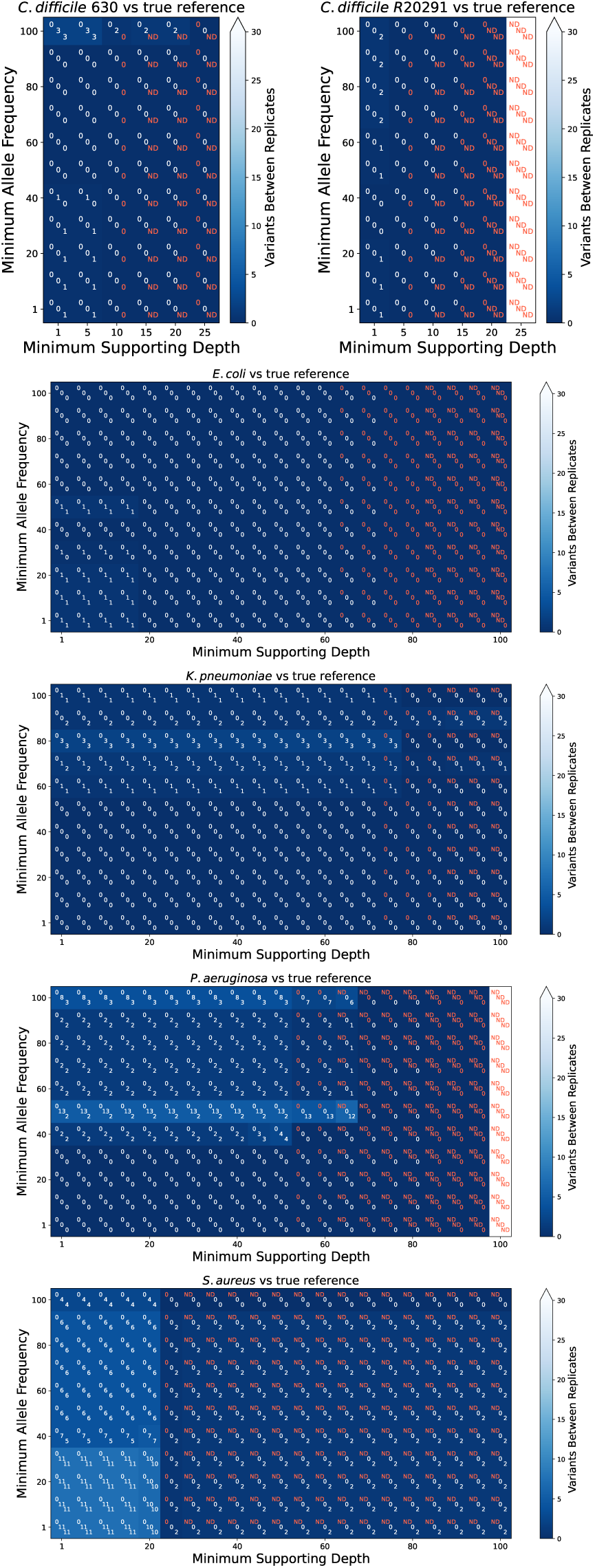
Number of false positives compared with the true reference, in the reference replicate set, by minimum frequency and minimum read depth supporting each call as reported by Clair3. The number of false positives for each replicate is shown in the centre of each cell: cells are coloured based on the number of false positives found with darker tones representing fewer variants (theoretical minimum 0). Due to the loss of breadth of coverage resulting from increasing the supporting depth, red text shows the number of variants found in combinations where less than 75% of the reference genome was called, suggesting that the fewer variants detected is likely caused by loss of data rather than optimal combination of variant frequency and supporting depths. Samples with less than the required depth are indicated by ND (no data). Note that higher stringency in the requirement of minimum supporting frequency and depth does not result in fewer false positives between replicates as Figure 4 illustrates with bcftools, but often more, with no clear relationship between number of false positives, minimum frequency, and minimum depth.

**Figure S8.**
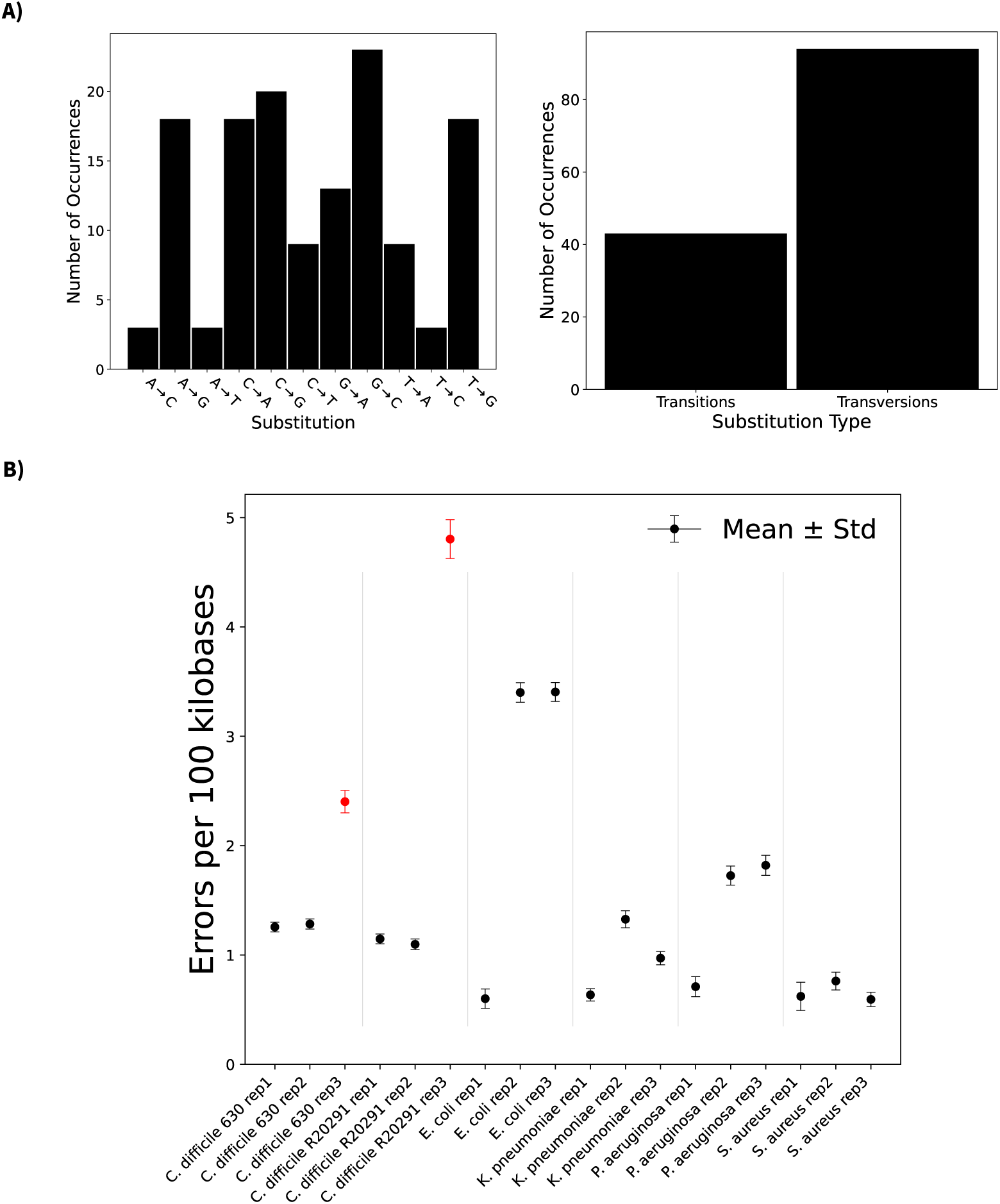
False positive variant analysis (A) and self-reported sequencing errors (B) in the replicate reference set when mapped to the true reference. **A)** Substitutions (REFERENCE→SAMPLE) using Dorado version 1.0.2 with super accuracy model (left) and whether these mutations correspond to transitions or transversions (right). **B)** Error per 100 kilobases (Kb) based on self-reported Phred read quality scores in Dorado version 1.0.2. We transformed the quality score using the expression^59^ 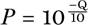 where Q is the mean Phred quality score of the read and *P* the probability of error. We then used this probability and the read length to estimate the error per nucleotide, and multiplied by 100,000. Replicates in red were excluded by ‘*OxBreaker*’ from downstream analysis due to coverage of the called genome being below 75%.

**Figure S9.**
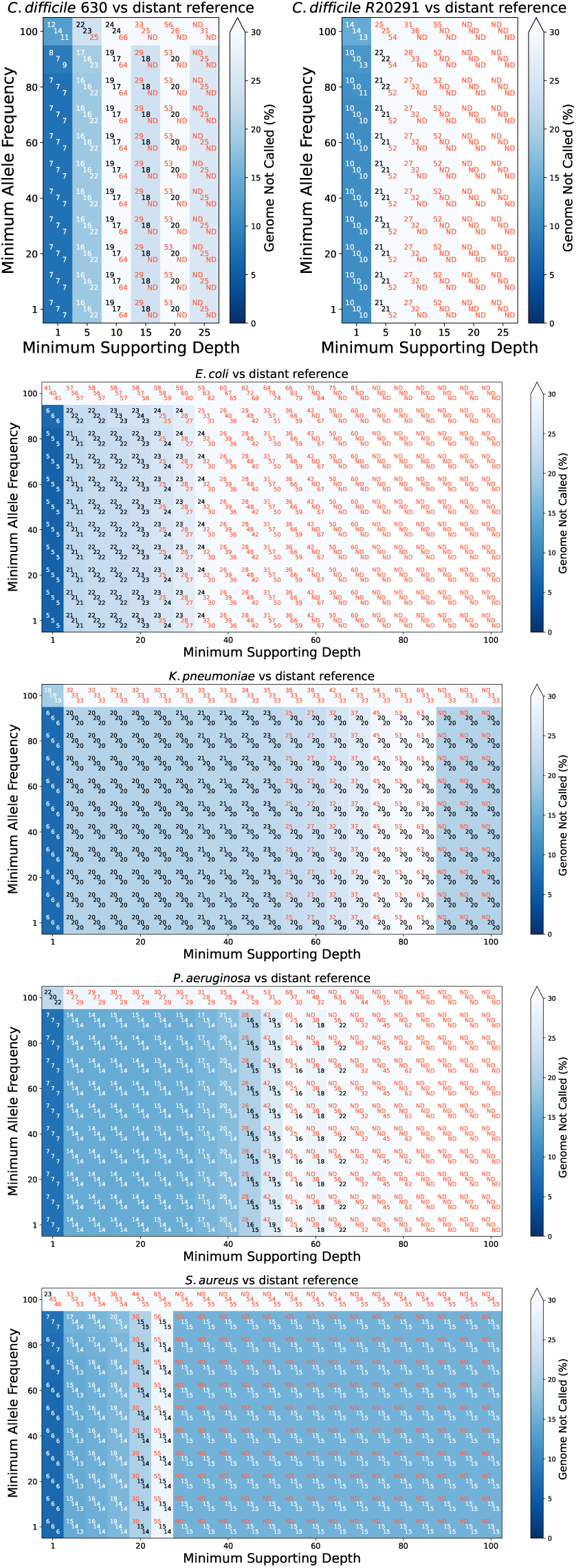
Percentage of genome not called in the replicate reference set when mapped against a distant reference genome. The percentage of genome not called for each replicate is shown in the centre of each cell, and cells are coloured when the percentage is above 25%, those with most percentage of their genome available for analysis being represented with darker tones (theoretical minimum 0). Samples with less than the required depth are noted as ND (no data).

**Figure S10.**
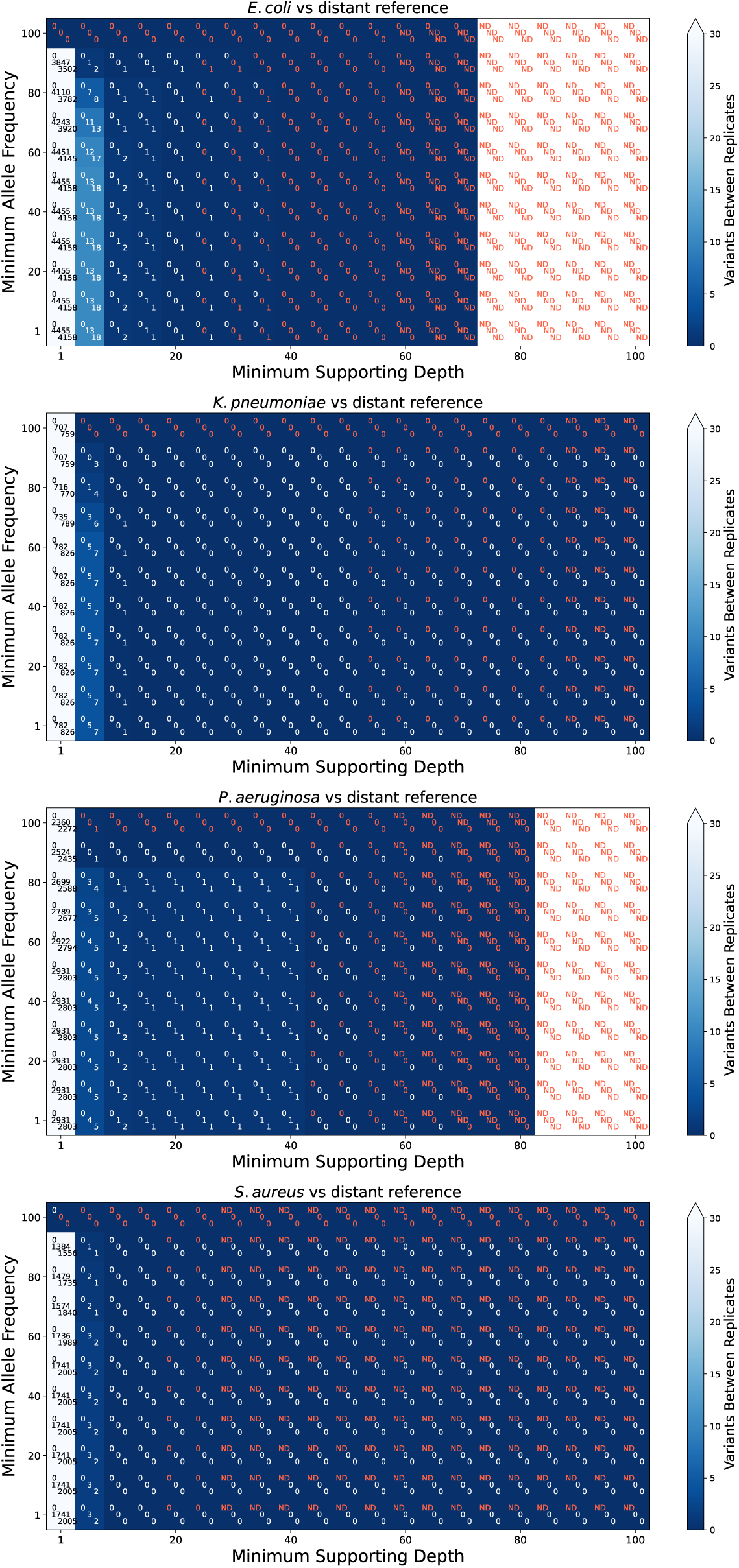
Number of false positives when mapping against a distant reference in replicate reference set isolates with depth >50X optimise minimum frequency and minimum read depth. The number of variants for each replicate is shown in the centre of each cell, and cells are coloured based on the number of variants found with darker tones representing fewer variants (theoretical minimum 0). Due to the loss of breadth of coverage resulting from increasing the supporting depth (Figure S2), red text denotes the number of variants found in combinations where less than 75% of the reference genome was called. Samples without depth data available are noted as ND (no data). This serves as a means to warn that the fewer variants is likely caused by loss of data rather than optimal combination of variant frequency and supporting depths.

## REFERENCES

1. Point prevalence survey on healthcare-associated infections, antimicrobial use and antimicrobial stewardship in England, 2023 tech. rep. (UK Health Security Agency (UKHSA), 2025).

2. Guest, J. F., Keating, T., Gould, D. & Wigglesworth, N. Modelling the annual NHS costs and outcomes attributable to healthcare-associated infections in England. BMJ open 10, e033367 (2020).

3. Sundermann, A. J., Kumar, P., Griffith, M. P., et al. Real-Time Genomic Surveillance for Enhanced Healthcare Outbreak Detection and Control: Clinical and Economic Impact. Clin. Infect. Dis., ciaf216 (2025).

4. Seth-Smith, H., Roloff, T., Benvenga, V. & Egli, A. Usage of bacterial whole genome sequencing: outbreaks and beyond in pediatric patients. The Pediatric Infectious Disease Journal 44, e53–e55 (2025).

5. Sundermann, A. J., Rosa, R., Harris, P. N., et al. Pathogen genomics in healthcare: overcoming barriers to proactive surveillance. Antimicrob. Agents Chemother. 69, e01479–24 (2025).

6. Goldin, M. R., Ruderfer, D. M., Bick, A., Roden, D. M., Schuler, B. A. & Robinson, J. R. Benefits and barriers to broad implementation of genomic sequencing in the NICU. The American Journal of Human Genetics (2025).

7. Greninger, A. L. & Zerr, D. M. NGSocomial infections: High-resolution views of hospital-acquired infections through genomic epidemiology. Journal of the Pediatric Infectious Diseases Society 10, S88–S95 (2021).

8. Pankhurst, L. J., del Ojo Elias, C., Votintseva, A. A., et al. Rapid, comprehensive, and affordable mycobacterial diagnosis with whole-genome sequencing: a prospective study. Lancet Respir. Med. 4, 49–58 (2016).

9. Deakin, C. T., Deakin, J. J., Ginn, S. L., et al. Impact of next-generation sequencing error on analysis of barcoded plasmid libraries of known complexity and sequence. Nucleic acids research 42, e129–e129 (2014).

10. Dimitriu, T. Evolution of horizontal transmission in antimicrobial resistance plasmids. Microbiology 168, 001214 (2022).

11. McInerney, J. O., McNally, A. & O’connell, M. J. Why prokaryotes have pangenomes. Nat. Microbiol 2, 1–5 (2017).

12. Valiente-Mullor, C., Beamud, B., Ansari, I., et al. One is not enough: on the effects of reference genome for the mapping and subsequent analyses of short-reads. PLoS Comput. Biol. 17, e1008678 (2021).

13. Pearce, M. E., Alikhan, N.-F., Dallman, T. J., Zhou, Z., Grant, K. & Maiden, M. C. Comparative analysis of core genome MLST and SNP typing within a European Salmonella serovar Enteritidis outbreak. Int. J. Food Microbiol. 274, 1–11 (2018).

14. Boers, S. A., Van der Reijden, W. A. & Jansen, R. High-throughput multilocus sequence typing: bringing molecular typing to the next level. PloS One 7, e39630 (2012).

15. Jackson, B. R., Tarr, C., Strain, E., et al. Implementation of nationwide real-time whole-genome sequencing to enhance listeriosis outbreak detection and investigation. Rev. Infect Dis. 63, 380–386 (2016).

16. Romano-Bertrand, S, Virieux-Petit, M, Mauffrey, F, Senn, L & Blanc, D. Defining a genomic threshold for investigating Pseudomonas aeruginosa hospital outbreak. J. Hosp. Infect. 161, 119–129 (2025).

17. Wang, Y., Zhao, Y., Bollas, A., Wang, Y. & Au, K. F. Nanopore sequencing technology, bioinformatics and applications. Nat. Biotechnol. 39, 1348–1365 (2021).

18. Cao, M. D., Ganesamoorthy, D., Elliott, A. G., Zhang, H., Cooper, M. A. & Coin, L. J. Streaming algorithms for identification pathogens and antibiotic resistance potential from realtime MinION™ sequencing. Gigascience 5, s13742–016 (2016).

19. Wagner, G. E., Dabernig-Heinz, J., Lipp, M., et al. Real-time nanopore Q20+ sequencing enables extremely fast and accurate core genome MLST typing and democratizes access to high-resolution bacterial pathogen surveillance. J. Clin. Microbiol. 61, e01631–22 (2023).

20. Bloomfield, M., Bakker, S., Burton, M., et al. The need for speed: ultra-rapid high-resolution outbreak analysis in a front-line hospital microbiology laboratory. J. Hosp. Infect. 168, 48–57. 10.1016/j.jhin.2025.11.020 (2026).

21. Iribarren, S. J., Sward, K. A., Beck, S. L., Pearce, P. F., Thurston, D. & Chirico, C. Qualitative evaluation of a text messaging intervention to support patients with active tuberculosis: implementation considerations. JMIR mHealth and uHealth 3, e3971 (2015).

22. Wood, D. E., Lu, J. & Langmead, B. Improved metagenomic analysis with Kraken 2. Genome biol. 20, 1–13 (2019).

23. Loman, N. J. Kraken 2 Microbial Database https://lomanlab.github.io/mockcommunity/mc_databases.html. 2018.

24. Tatusova, T., Ciufo, S., Fedorov, B., O’Neill, K. & Tolstoy, I. RefSeq microbial genomes database: new representation and annotation strategy. Nucleic Acids Res. 42, D553–D559 (2013).

25. Cock, P. J., Antao, T., Chang, J. T., et al. Biopython: freely available Python tools for computational molecular biology and bioinformatics. Bioinformatics 25, 1422 (2009).

26. Liu-Wei, W., van der Toorn, W., Bohn, P., Hölzer, M., Smyth, R. P. & von Kleist, M. Sequencing accuracy and systematic errors of nanopore direct RNA sequencing. BMC Genomics 25, 528 (2024).

27. Ye, F., Zhu, J., Zhang, X., et al. Characteristics and filtering of low-frequency artificial short deletion variations based on nanopore sequencing. GigaScience 14, giaf018 (2025).

28. Thomas, C., Brangsch, H., Galeone, V., Hölzer, M., Marz, M. & Linde, J. Accurately assembling nanopore sequencing data of highly pathogenic bacteria. BMC Genomics 26, 783 (2025).

29. Camacho, C., Coulouris, G., Avagyan, V., et al. BLAST+: architecture and applications. BMC Bioinformatics 10, 1–9 (2009).

30. Lerat, E. Identifying repeats and transposable elements in sequenced genomes: how to find your way through the dense forest of programs. Heredity 104, 520–533 (2010).

31. Delihas, N. Impact of small repeat sequences on bacterial genome evolution. Genome Biol. Evol. 3, 959–973 (2011).

32. Li, H. Minimap2: pairwise alignment for nucleotide sequences. Bioinformatics 34, 3094– 3100 (2018).

33. Li, H., Handsaker, B., Wysoker, A., et al. The sequence alignment/map format and SAM-tools. Bioinformatics 25, 2078–2079 (2009).

34. Seemann, T. Source code for snp-dists software. Zenodo (2018).

35. Croucher, N. J., Page, A. J., Connor, T. R., et al. Rapid phylogenetic analysis of large samples of recombinant bacterial whole genome sequences using Gubbins. Nucl. Acids Res. 43, e15– e15 (2015).

36. Eyre, D. W., Cule, M. L., Wilson, D. J., et al. Diverse sources of C. difficile infection identified on whole-genome sequencing. N. Eng. J. Med. 369, 1195–1205 (2013).

37. Hall, M. D., Holden, M. T., Srisomang, P., et al. Improved characterisation of MRSA transmission using within-host bacterial sequence diversity. eLife 8, e46402 (2019).

38. Coll, F., Raven, K. E., Knight, G. M., et al. Definition of a genetic relatedness cutoff to exclude recent transmission of meticillin-resistant Staphylococcus aureus: a genomic epidemiology analysis. Lancet Microbe 1, e328–e335 (2020).

39. Dabernig-Heinz, J., Lohde, M., Hölzer, M., et al. A multicenter study on accuracy and reproducibility of nanopore sequencing-based genotyping of bacterial pathogens. J. Clin. Microbiol. 62, e00628–24 (2024).

40. Zheng, Z., Li, S., Su, J., Leung, A. W.-S., Lam, T.-W. & Luo, R. Symphonizing pileup and full-alignment for deep learning-based long-read variant calling. Nat. Comput. Sci. 2, 797– 803 (2022).

41. Hall, M. B., Wick, R. R., Judd, L. M., et al. Benchmarking reveals superiority of deep learning variant callers on bacterial nanopore sequence data. eLife 13, RP98300 (2024).

42. Bogaerts, B., Maex, M., Commans, F., et al. Oxford Nanopore Technologies R10 sequencing enables accurate cgMLST-based bacterial outbreak investigation of Neisseria meningitidis and Salmonella enterica when accounting for methylation-related errors. J. Clin. Microbiol. 63, e00410–25 (2025).

43. Sanderson, N. D., Hopkins, K. M., Colpus, M., et al. Evaluation of the accuracy of bacterial genome reconstruction with Oxford Nanopore R10. 4.1 long-read-only sequencing. Microb. Genom. 10, 001246 (2024).

44. Lees, E. A., Gentry, J., Webster, H., et al. Multiple introductions of NRCS-A Staphylococcus capitis to the neonatal intensive care unit drive neonatal bloodstream infections: a case-control and environmental genomic survey. Microb. Genom. 11, 001340 (2025).

45. Moran, R. A., Behruznia, M., Holden, E., Garvey, M. I. & McNally, A. pQEB1: A hospital outbreak plasmid lineage carrying bla KPC-2. Microb. Genom. 10, 001291 (2024).

46. Bush, S. J., Foster, D., Eyre, D. W., et al. Genomic diversity affects the accuracy of bacterial single-nucleotide polymorphism–calling pipelines. GigaScience 9, giaa007 (2020).

47. Shibata, Y., Tien, L. H. T., Nomoto, R. & Osawa, R. Development of a multilocus sequence typing scheme for Streptococcus gallolyticus. Microbiology 160, 113–122 (2014).

48. Delahaye, C. & Nicolas, J. Sequencing DNA with nanopores: Troubles and biases. PloS ONE 16, e0257521 (2021).

49. Coll, F. Key variables affecting genetic distance calculations in genomic epidemiology. Lancet Microbe 2, e486–e487 (2021).

50. Gorrie, C. L., Da Silva, A. G., Ingle, D. J., et al. Key parameters for genomics-based real-time detection and tracking of multidrug-resistant bacteria: a systematic analysis. The Lancet Microbe 2, e575–e583 (2021).

51. Van der Roest, B. R., Bootsma, M. C., Fischer, E. A., et al. Phylodynamic assessment of SNP distances from whole genome sequencing for determining Mycobacterium tuberculosis transmission. Sci. Rep. 15, 10694 (2025).

52. Derelle, R., von Wachsmann, J., Mäklin, T., et al. Seamless, rapid, and accurate analyses of outbreak genomic data using split k-mer analysis. Genome Res. 34, 1661–1673 (2024).

53. Vereecke, N., Yoon, T. B., Luo, T. L., et al. An open-source nanopore-only sequencing workflow for analysis of clonal outbreaks delivers short-read level accuracy. J. Clin. Microbiol. 63, e00664–25 (2025).

54. Cottingham, H., Judd, L. M., Harshegyi-Hand, T., et al. Nanopore sequencing enables highly accurate genotyping and identification of resistance determinants in key nosocomial pathogens. medRxiv, 2025–07 (2025).

55. Van Boven, L. S., Kusters, R. W., Klokman, V. W., Dameff, C. & Barten, D. G. Acute care disruptions due to information technology failures in the Netherlands from 2000 to 2020. Health Policy and Technology 13, 100840 (2024).

56. Tully, J. L., Rao, S., Straw, I., et al. Patient care technology disruptions associated with the CrowdStrike outage. JAMA Network Open 8, e2530226 (2025).

57. Hertz, F. B., Nielsen, K. L., Strunin, D., et al. Estimating the potential economic and health impact of integrated genomic surveillance in a hospital setting. Clin. Microbiol. Infect. (2025).

58. De Coster, W. & Rademakers, R. NanoPack2: population-scale evaluation of long-read sequencing data. Bioinformatics 39, btad311 (2023).

59. Ewing, B. & Green, P. Base-calling of automated sequencer traces using phred. II. Error probabilities. Genome Res. 8, 186–194 (1998).

